# Allele frequency dynamics under sex-biased demography and sex-specific inheritance in a pedigreed jay population

**DOI:** 10.1101/2021.10.28.466320

**Authors:** Rose M.H. Driscoll, Felix E.G. Beaudry, Elissa J. Cosgrove, John W. Fitzpatrick, Stephan J. Schoech, Nancy Chen

## Abstract

Sex-biased demography, including sex-biased survival or migration, can alter allele frequency changes across the genome. In particular, we can expect different patterns of genetic variation on autosomes and sex chromosomes due to sex-specific differences in life histories, as well as differences in effective population size, transmission modes, and the strength and mode of selection. Here, we demonstrate the role that sex differences in life history played in shaping short-term evolutionary dynamics across the genome. We used a 25-year pedigree and genomic dataset from a long-studied population of Florida Scrub-Jays (*Aphelocoma coerulescens*) to directly characterize the relative roles of sex-biased demography and inheritance in shaping genome-wide allele frequency trajectories. We used gene dropping simulations to estimate individual genetic contributions to future generations and to model drift and immigration on the known pedigree. We quantified differential expected genetic contributions of males and females over time, showing the impact of sex-biased dispersal in a monogamous system. Due to female-biased dispersal, more autosomal variation is introduced by female immigrants. However, due to male-biased transmission, more Z variation is introduced by male immigrants. Finally, we partitioned the proportion of variance in allele frequency change through time due to male and female contributions. Overall, most allele frequency change is due to variance in survival and births. Males and females make similar contributions to autosomal allele frequency change, but males make higher contributions to allele frequency change on the Z chromosome. Our work shows the importance of understanding sex-specific demographic processes in characterizing genome-wide allele frequency change in wild populations.

## Introduction

A fundamental goal of evolutionary biology is to determine the roles of different evolutionary processes in governing allele frequency change over time. Understanding evolution over contemporary timescales is especially important given its relevance to current issues such as public health (1), conservation policy (2, 3), and agricultural practices (4). To date, most population genetic studies use allele frequencies estimated from present-day samples to make inferences about the evolutionary processes that generated the observed patterns of genetic variation (5, 6), though studies that directly track allele frequencies over time are becoming more common (*e*.*g*., 7, 8). Building from a rich literature regarding evolution on ecological time scales (*e*.*g*., 9–11), recent studiesare adding to our understanding of the drivers of allele frequency change over short timescales in natural populations (12–15). Temporal genomic data coupled with knowledge of the population pedigree, or the relationships among all individuals in a population over time, can provide precise estimates of the mechanisms underlying short-term allele frequency dynamics (16).

Allele frequency change is driven by differential survival, reproduction, and dispersal among individuals. In other words, depending on their life history, different individuals have different genetic contributions to a population over time (17). The expected genetic contribution of an individual is defined as the expected number of alleles inherited from that individual present in the population in future generations, and is determined by both the number of descendants of an individual and the randomness of chromosomal segregation and recombination (18, 19). Individual expected genetic contributions are expected to stabilize after a few generations and can be used to estimate individual reproductive values (6, 18, 19).

One important factor affecting an individual’s genetic contribution over time is its sex. In sexually-reproducing organisms, the expected genetic contributions of males and females should be equal on average (6). However, different sexes often have different life history traits: depending on their mating system, individuals of different sexes can differ in the variance in number of offspring (*i*.*e*., reproductive success; 20, 21), dispersal likelihood and distance (22), or niche occupation (23). These differences in life history traits can lead to population-level effects such as a biased adult sex ratio or sex-biased migration rates (24–26). Differences in life history between individuals of different sexes may affect expected genetic contributions and therefore allele frequency change. Indeed, including sex increases the accuracy of modelling demographic changes (27, 28) and will likely help the characterization of sources of allele frequency change over time.

Individual expected genetic contributions can vary across the genome. For example, sex chromosomes should have different patterns of expected genetic contributions compared to autosomes. The two most common sex chromosome systems include X and Y chromosomes, where females are XX and males XY (*e*.*g*., in mammals, beetles, and many dipterans), and Z and W chromosomes, where females are ZW and males ZZ (*e*.*g*., in birds, lepidopterans, and strawberries). As a result of sex-specific differences in transmission (inheritance rules) and ploidy (*e*.*g*., hemizygosity), expected genetic contributions on sex chromosomes are disproportionately influenced by sex-biased demography. The X and Z chromosomes can have different effective population sizes than autosomes and are expected to experience more drift than the autosomes (29, 30). Indeed, different patterns of genetic variation on autosomes and sex chromosomes are expected to reflect sex-specific differences in effective population size and transmission modes, as well as in life histories (25, 30–34).

Based on differences in effective population size between autosomes and sex chromosomes, two expectations arise in ZW systems (35–37). First is the well-known prediction that males are expected to contribute 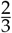 of Z variation, while females are expected to contribute 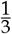, based on the number of chromosomes in each sex (25, 38). Second, the ratio of Z to autosome diversity should equal 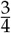 (39). These expectations require many simplifying assumptions, including statistical equilibrium as well as equal mutation rates and fitness variance between the sexes (39). Departures from these expectations can result from sex-biased demographic processes, fuelling the use of neutral genetic diversity ratios between autosomes and sex chromosomes to infer sex-biased demography over long timescales, (*i*.*e*., ≫ Ne generations). For example, the strength of sexual selection, defined as sex-biased variance in reproductive success (40), can be estimated from genetic diversity differences between X and autosomes (29, 41–43). Similarly, sex-biased migration can be inferred from sex chromosome versus autosome neutral genetic diversity ratios (31, 44).

These endeavours, however, have important caveats. First, it is challenging (if not impossible) to disentangle which sex-biased processes are contributing to biased genetic diversity ratios when several processes occur jointly, especially in populations with overlapping generations of variable sizes (29). Second, for inferences of sex-biased migration, timescale has been shown to have important effects on genetic diversity ratios in humans (45, 46), but timescale is rarely considered in inferences of gene flow (47). Third, patterns of neutral genetic diversity are affected by linked selection and therefore vary depending on differences in effective recombination rate between sex chromosomes and autosomes as well as between the sexes (30). These caveats suggest that approaches complementary to neutral genetic diversity data are necessary to deepen our understanding of the effects of sex-biased demography in natural populations.

One approach for studying sex-biased demography is to use individual genetic contributions. Expected genetic contributions can be estimated from population pedigrees using analytical calculations (48, 49) or simulations on the pedigree (gene dropping; (50)). With the advent of high-throughput molecular sequencing, gene dropping has been adopted to simulate changes in allele frequency based on known genotypes. This method has been used to estimate the frequency of known recessive lethals in cattle (51), humans (52) and Soay sheep (53), as well as to identify alleles thought to be under positive selection in Florida Scrub-Jays (16), Soay Sheep (54) and domestic foxes (55). Population pedigrees are therefore an invaluable resource for studying the impacts of sex-biased demography on allele frequency change over time.

A recent study characterized short-term allele frequency dynamics at autosomal loci in a population of Federally Threatened Florida Scrub-Jays (*Aphelocoma coerulescens*) by combining simulations of expected genetic contributions with genomic data (16). Florida Scrub-Jays are cooperatively breeding birds with a balanced sex ratio of breeding adults, as these jays are socially and genetically monogamous (56, 57). Natal dispersal is limited and female-biased (58–60). Previous work found that immigration plays a large role in allele frequency change even in a large population of jays (16, 61). A population of jays at Archbold Biological Station (hereafter Archbold) has been closely monitored for over 50 years, resulting in a detailed population pedigree with over 6,000 individuals. This study population provides an unrivalled opportunity to study individual-level effects on short-term allele frequency change.

Here, we extended the work of (16) by developing and applying gene dropping methods for sex chromosomes to directly evaluate the contributions of sex-biased demographic processes (*i*.*e*., dispersal) and sex-specific inheritance to changes in allele frequencies over time in the Florida Scrub-Jay. We first estimated the expected genetic contributions of individuals, contrasting the role of males and females as well as of immigrants grouped by sex. Using separate models to track different modes of inheritance, we compared the expected genetic contributions of males and females at autosomal and Z-linked loci. Next, to identify loci likely to be under selection, we incorporated genotype information and simulated changes in allele frequency over time given the pedigree. Finally, we partitioned the variance in allele frequency change between years across sexes and demographic groups, and quantified the contributions of males and females and different evolutionary processes to allele frequency change at autosomal and Z-linked loci.

## Methods

### Study population and genomic data

A population of Florida Scrub-Jays at Archbold in Venus, Florida, USA, has been intensively monitored for multiple decades, the northern half since 1969 (56) and the southern half since 1989 (62, 63). Each bird in this population has been uniquely banded, making immigrants easy to identify. Immigrants are defined as birds born outside the study population, but the source population of any given immigrant is not known. The entire population is surveyed every few months to provide detailed records of individual survival. Family groups are closely monitored to assess reproductive success, with new nestlings officially banded at 11 days of age. As a result of these detailed records, we have a fairly comprehensive pedigree of the entire population. Pedigree relationships based on field observations are mostly accurate because of the low rate of extra-pair paternity in this species (57), and we confirmed pedigree relationships of >3,000 individuals with genomic data (61). To avoid the complications associated with study tract expansion in the 1980s, we truncated the pedigree at 1990, resulting in a pedigree with 6,936 individuals in total by the end of our study period in 2013. We note that there was a minor study tract expansion in the southern end of our field site in 1993 that partially contributed to the observed peak in immigration in 1994 (as we cannot distinguish true immigrants from previously-unbanded residents in the new study tract area); however, we also observed an increase in known immigrants in the northern half of our field site (a consistently monitored area) in 1994. The addition of these territories did not impact overall immigration dynamics. Fieldwork at Archbold Biological Station was approved by the Institutional Animal Care and Use Committees at Cornell University (IACUC 2010-0015), the University of Memphis (0067), the University of Rochester (102153), and Archbold Biological Station (AUP-006-R), and was permitted by the U.S. Fish and Wildlife Service (TE824723-8, TE-117769), the U.S. Geological Survey (banding permits: 07732, 23098), and the Florida Fish and Wildlife Conservation Commission (LSSC-10-00205).

Since 1999, birds in the Archbold population have been sexed using molecular markers, following the protocol of Fridolfsson and Ellegren (64). A previous study generated genotype data for 3,984 individuals using a custom Illumina iSelect Beadchip containing 15,416 single nucleotide polymorphisms (SNPs) (61). These individuals were from a set of 68 territories that were consistently monitored since 1990, and included near-exhaustive sampling in 1989-1991, 1995, and 1999-2013. Here, we used data from a set of 10,731 autosomal SNPs previously used in (16) (see 16, 61, for more information on SNP discovery, filtering, and quality control) along with 250 Z-linked and 19 pseudoautosomal SNPs with minor allele frequency > 0.05. We performed additional scaffolding for the Florida Scrub-Jay genome (NCBI BioProject PRJNA1076903) and annotated the genome and our SNPs (see Supplementary Material). In this version of the genome assembly, the Z chromosome represents 7.1% of the total genome (75.6 Mb of 1.06 Gb total; Table S1).

### Expected genetic contributions of males and females

We begin by calculating the genealogical contribution of each individual, defined as the proportion of a given birth cohort that is descended from that individual. In contrast, we estimated the expected genetic contribution of an individual as the expected proportion of alleles in a given birth cohort that have been transmitted from our focal individual. Expected individual genetic contributions depend on transmission patterns and therefore should differ for autosomal versus sex-linked loci.

For a given autosomal locus, fathers and mothers each transmit one of their two alleles randomly to each offspring. Thus, the expected genetic contribution of an individual may be obtained by tracing the transmission of their alleles through the pedigree. The expected genetic contribution of an individual at an autosomal locus (*G*_*auto*_) is:

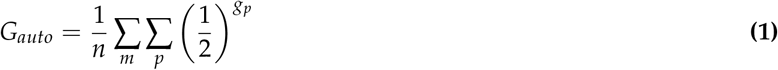

where *n* is the number of nestlings in the birth cohort, *m* is the number of nestlings in the cohort related to the focal individual (their descendants), *p* is the number of paths in the pedigree linking the focal individual and a given descendant, and *g*_*p*_ is the number of generations between the focal individual and the descendant along a given path.

For a given Z chromosome locus, fathers transmit one of their two Z chromosomes randomly to each offspring, but mothers invariably transmit their single Z chromosome to their male offspring only. The expected genetic contribution of an individual at a Z chromosome locus in a given year (*G*_*Z*_) is:

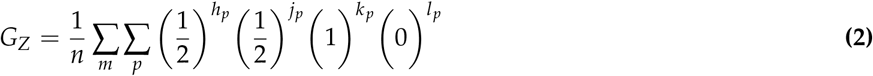

where *n, m*, and *p* are as described above. *h*_*p*_ represents the number of male-to-male (*i*.*e*., father-to-son) transmission events, *j*_*p*_ represents the number of male-to-female (*i*.*e*., father-to-daughter) transmission events, *k*_*p*_ represents the number of female-to-male (*i*.*e*., mother-to-son) transmission events, and *l*_*p*_ represents the number of female-to-female (*i*.*e*., mother-to-daughter) transmission events in a given path. Note that a female-to-female transmission anywhere in a path will always result in zero genetic contribution from the focal individual to the descendant along that path, as females never pass on their Z to their daughters. For a given path from a focal individual to a descendant, *h*_*p*_ + *j*_*p*_ + *k*_*p*_ + *l*_*p*_ = *g*_*p*_. An equivalent way of estimating expected genetic contributions is to simulate the transfer of alleles down the pedigree using gene dropping, emulating random segregation at meiosis (16). This method is preferable because it allows us to determine the variance across loci around a calculated expectation, and so we use the simulation method here.

To assess the expected autosomal genetic contributions of a single focal individual, we truncated the pedigree above the focal individual to make them a “founder”, *i*.*e*., an individual with no known parents in the pedigree. We then assigned the focal individual a genotype of ‘22’ and all other founders a genotype of ‘11’, simulated Mendelian transmission of alleles down the pedigree, and calculated the frequency of the ‘2’ allele in year *t*. We repeated this process for one million iterations. To assess expected Z chromosome genetic contributions, we used a similar method, but adjusted transmission rules and assigned females a genotype of ‘2’ (the focal individual) or ‘1’ (all other founders) for their single copy of the Z chromosome. In this section, we assign individuals genotypes instead of using observed genotype data because we want to quantify the expected genetic contributions of specific individuals, and simulations using observed genotypes would confound the contributions of all individuals who share an allele at that locus. To perform these simulations, we needed to know the sex of all descendants. For the 1,896 unsexed individuals in our population pedigree (all died as nestlings), we assigned individuals with missing sex information as male; since males are ZZ, this approach will minimize Z-autosome differences. Note that since the sex of all parents is known and only 1.4% of nestlings born after 2005 have missing sex information, how we deal with unsexed individuals does not significantly change our results (Supplementary Material; Fig. S1).

We estimated the genealogical and mean expected genetic contributions over time for a set of 926 breeders that bred in the population between 1990 and 2013. These individuals were all born before 2002 and died by the end of 2014, so our estimates should capture all of their offspring as well as some grand-offspring, great-grand-offspring, etc. To test the relationship between an individual’s sex and expected genetic contribution to the population in 2013, we fit linear models for expected genetic contributions with genealogical contribution, sex, and their interaction as independent variables. We next fit linear models for the ratio of Z/autosome expected genetic contributions using the expected genetic contribution on the Z chromosome as the dependent variable and autosomal expected contribution as the independent variable. We also modeled expected genetic contributions on the Z with autosomal expected contribution, sex, and their interaction as independent variables.

### Expected genetic contributions of male and female immigrants

To quantify the effects of sex-biased migration, we estimated the expected genetic contributions of male and female immigrants for both autosomal and Z-linked loci. Starting with immigrants entering the population in 1991, we assigned all male immigrants a genotype of ‘22’ and all other founders a genotype of ‘11’, performed gene dropping using the appropriate transmission rules as described above, and calculated the frequency of the ‘2’ allele in each year. We assigned individuals with missing sex information as male, as above. We repeated this process one million times to find the total expected genetic contributions of all male immigrants as a group. We repeated this process for female immigrants. Note that the total expected genetic contribution of immigrants here is cumulative and includes the contributions of all descendants of immigrants. We used linear models to test for trends in immigration rate and sex ratios of immigrants over time. We included the number of immigrants in our model of sex ratios. Finally, we used linear models to assess the relationship between immigrant cohort size and the expected genetic contribution of that immigrant cohort to the population 15 years later (the longest time period for which we have a large enough data set).

### Neutral allele dynamics and selection

Previous work evaluated signals of selection on 10,731 autosomal loci and detected 18 SNPs with significant changes in allele frequency across 1999-2013 (16). Here, we tested for selection on the Z chromosome using 250 Z-linked SNPs and 19 pseudoautosomal SNPs. Methods and results for the pseudoautosomal region can be found in Supplementary Material. To simulate the neutral behavior of Z-linked alleles, we performed gene dropping simulations for each SNP using observed founder genotypes. We trimmed the 6,936-individual pedigree to remove any founders with missing genotype data for the focal SNP. We re-coded the offspring of trimmed founders as founders, and repeated this process until all founders had genotype data. Thus, the pedigree used for each SNP could vary slightly based on patterns of missing data at that locus. After trimming, we used gene dropping to simulate the transmission of alleles down the pedigree one million times, using Z chromosome transmission rules. From the simulated genotypes in each iteration, we calculated the expected allele frequency shifts using the genotyped nestlings born in each year in a core set of about 68 consistently monitored territories (2,841 nestlings in total over 1990-2013). We randomly assigned unsexed individuals as males or females based on the empirical sex ratio (*∼*50:50), but results are similar even if we assign all unsexed individuals as males because no unsexed individuals are genotyped or produced offspring. These gene dropping iterations resulted in a distribution of changes in allele frequency over time, conditional on the pedigree. We compared the distribution of allele frequency change expected under neutrality to observed allele frequency changes between 1999 and 2013 and in adjacent years during that time period. We determined p-values for allele frequency change at each SNP by counting the number of simulations in which the simulated value was further from the median of the expected distribution than the observed value. We performed false discovery rate (FDR) correction of p-values for all comparisons across both the autosomes (16) and the Z (including the 19 pseudoautosomal SNPs; 11,000 SNPs total) and applied a threshold of FDR < 0.1 or 0.25 to assess significance.

### Model of variance in allele frequencies

To determine the sources of genome-wide allele frequency change for autosomes and the Z chromosome, we extended the model from (16) to incorporate the sex of individuals. Like (16), we assumed only three sources of allele frequency change: differences in survival, reproduction (*i*.*e*., new births), and immigration among individuals. We classified individuals in our population census each year as survivors, immigrants, or nestlings/births. For this analysis (in contrast to previous analyses), immigrating individuals were only counted as immigrants the year they first appear in the population and were categorized as survivors in following years. We further subdivided each demographic group into males and females in order to test the possibility that individuals of different sexes contributed differently to each source of allele frequency change. We included individuals in the population for a given year if they were observed during at least two months between March and June, and considered individuals who disappeared from the population for one or more years but later returned to have been survivors in the intervening time.

We calculated allele frequency change between consecutive years by contrasting allele frequencies in each category (male survivors, male births, male immigrants, female survivors, female births, female immigrants) in each year with the allele frequency of the entire population in the previous year. Let *N*_*t*_ be the total number of individuals in the population in year *t*. Let *N*_*M,s*_ be the number of males who survived from year *t −* 1 to year *t, N*_*M,i*_ be the number of males who immigrated into the population in year *t*, and *N*_*M,b*_ be the number of males born in the population in year *t*. Likewise, let *N*_*F,s*_ be the number of females who survived from year *t −* 1 to year *t, N*_*F,i*_ be the number of females who immigrated into the population in year *t*, and *N*_*F,b*_ be the number of females born in the population in year *t*. Note that *N*_*t*_ = *N*_*M,s*_ + *N*_*M,i*_ + *N*_*M,b*_ + *N*_*F,s*_ + *N*_*F,i*_ + *N*_*F,b*_.

We constructed two models, one for autosomal loci and one for Z-linked loci. Let the allele frequency of a given demographic group *j* be *p*_*M,j*_ for males and *p*_*F,j*_ for females, *e*.*g*., *p*_*F,s*_ is the allele frequency of female survivors. The change in allele frequencies between two adjacent years for an autosomal locus is:

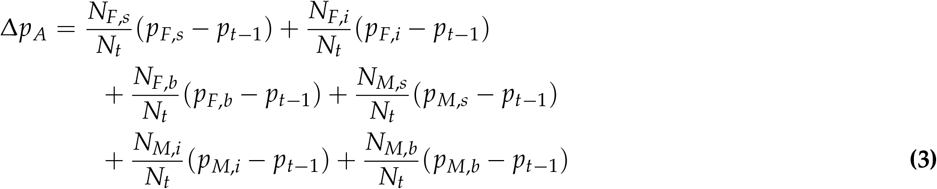

Here, the contribution of each category to the overall change in allele frequency over time is a function of the proportion of alleles in the population from that category in year *t* (equivalent to the proportion of individuals in that category) and the difference in allele frequencies between that category in year *t* (*e*.*g*., female survivors, *p*_*F,s*_) and the entire population from the year before (*p*_*t−*1_).

Similarly, the change in allele frequencies between two years for a Z-linked locus is:

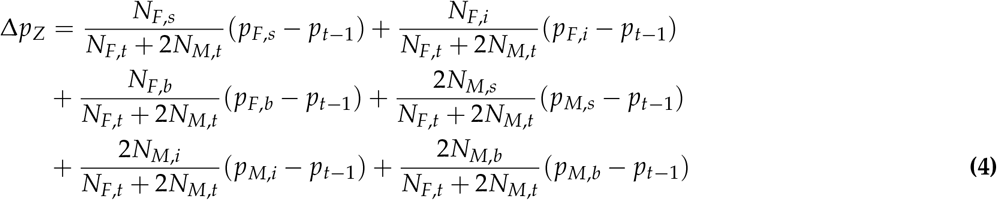

Note that males have two copies of the Z while females only have one copy of the Z, which affects the total number of alleles in the population and the relative weighting of male vs. female categories.

Using equations 3 and 4, we calculated the variance in allele frequency change between years for autosomal or Z-linked loci in each category, as well as the covariances between categories, *e*.*g*., between survival and birth (see Appendix A). We assumed that immigrants in a given year are unrelated to survivors and therefore set all covariances between immigrants and survivors to zero. In addition, for Z-linked loci, female survivors and female immigrants cannot transmit alleles to female nestlings (births), so we also set covariances between female survivors and female nestlings and between female immigrants and female nestlings to zero. We then calculated the proportion of total allele frequency change between a given pair of consecutive years contributed by each variance and covariance term.

For both models, we further partitioned allele frequency change associated with births, Var(*p*_*M,b*_ *− p*_*t−*1_) and Var(*p*_*F,b*_ *− p*_*t−*1_), into contributions from variation in family size and Mendelian segregation of alleles from heterozygous parents in order to understand the relative roles of these two factors (see full derivation in Appendix A).

While we have a complete census of all individuals in the population, not every individual is genotyped or sexed (Fig. S2). To deal with missing genotype data, we corrected each term for sampling error, estimated empirically via simulations. Briefly, we assigned genotypes to all individuals in the population in 1990 based on an allele frequency drawn from the empirical allele frequency distribution and then simulated Mendelian transmission (of autosomes or of the Z chromosome) forward in time for 100,000 loci. For our Z chromosome simulations, we randomly assigned unsexed individuals as males or females based on the empirical sex ratio, which is approximately 50:50. All unsexed individuals were also ungenotyped. We estimated sample allele frequencies from the subset of individuals who were genotyped and subtracted the population allele frequency to obtain the sampling error for our allele frequency estimates. We then estimated the errors for each term in our model using the equations in Appendix A.

To characterize the variance in allele frequency change over time in 1999-2013, we calculated allele frequencies among genotyped individuals for each category in each year at 10,731 autosomal SNPs for the autosomal model and 250 Z-linked SNPs for the Z chromosome model. Pseudoautosomal SNPs were not included in this analysis. For each model, we averaged across all loci to obtain the variance in allele frequency change contributed by each term (see equations 2 and 14 in Appendix A) and then corrected our estimates using our empirical estimates of sampling error (Supplementary Materials).

We used a bootstrapping approach to generate confidence intervals for our variance and covariance estimates. We sampled SNPs in 3.4 Mb windows along the genome 1,000 times with replacement. A 3.4 Mb window size was chosen to allow at least 1 window per chromosome, as the smallest chromosome in the current Florida Scrub-Jay genome assembly is 3.4 Mb. Windows smaller than 3.4 Mb, on unplaced scaffolds, or with less than 5 SNPs were discarded. Across autosomes, this windowed approach resulted in 299 windows containing between 5 and 257 SNPs, with 50 SNPs/window on average. On the Z, SNPs were split between 21 windows, ranging from 5-83 SNPs. To ensure that our chosen window size (3.4 Mb) was likely to break apart linkage, we estimated levels of linkage disequilibrium (LD) in our study population using the ‘ld-window-r2’ function in PLINK (65) to estimate pairwise LD between SNPs in each of our linkage groups. We fit a hyperbolic decay curve for the relationship between pairwise LD and distance between SNPs (as per 66) for autosomes and the Z chromosome separately and found that, on average, LD decayed around 0.1 Mb (Fig. S3). We repeated the above sampling and error estimation steps for each window. We also repeated this analysis with LD-pruned SNPs (Supplementary Material). For the full derivation of the autosomal model and for the model for Z-linked loci, see Appendix A.

### Implementation

We implemented gene dropping in python and C++ and used this software to conduct the simulations for expected genetic contributions and tests of selection. We performed analyses and visualization in R v. 3.6.3 (67) using packages base, stats, plyr (68), dplyr (69), ggplot2 (70), cowplot (71), and kinship2 (72).

We implemented the allele frequency variance models in R v. 4.1.1 (67) using packages base, stats, foreach (73), doParallel (74), plyr, and dplyr. We used the packages ggplot2 and cowplot for visualization.

### Data Availability

All scripts and data are available at https://github.com/felixbeaudry/ZDropping.

## Results

### Expected genetic contributions of males and females

We first focused on the contributions of individual breeding adults to future generations. Each individual breeder has both a genealogical and expected genetic contribution to the future. Here, we define the genealogical contribution of an individual as the proportion of a given birth cohort that is descended from that individual. In contrast, the expected genetic contribution of an individual is the expected proportion of alleles in a given birth cohort inherited identical-by-descent from our focal individual. An individual’s expected genetic contribution differs from their genealogical contribution because not every allele is passed down to the next generation, due to the randomness of Mendelian segregation. Expected genetic contributions are also affected by transmission rules; sex-biased transmission of the Z chromosome results in different genetic contributions of individuals at autosomal versus Z loci.

To illustrate the effect of sex-biased transmission on individual genetic contributions across the genome, we contrasted the expected genetic contributions for autosomal and Z-linked loci over time for a male and female who are each other’s exclusive mates. This pair first bred in 2001, produced 15 offspring together, and had a total of 223 descendants by 2013 (Fig. 1A). Since these breeders did not pair with any other individuals over the course of their lives, they have the same descendants and thus equal genealogical contributions. At a given autosomal locus, the male and female had equal expected genetic contributions (Fig 1B). The only differences in the expected autosomal genetic contributions of the male and female arose from the stochastic nature of the simulations we used to estimate expected genetic contributions; after 1,000,000 iterations, their values were essentially identical. In contrast, the expected genetic contributions on the Z chromosome tended to be considerably higher for the male than for the female between 2005 and 2013 (Fig. 1C).

**Figure 1.**
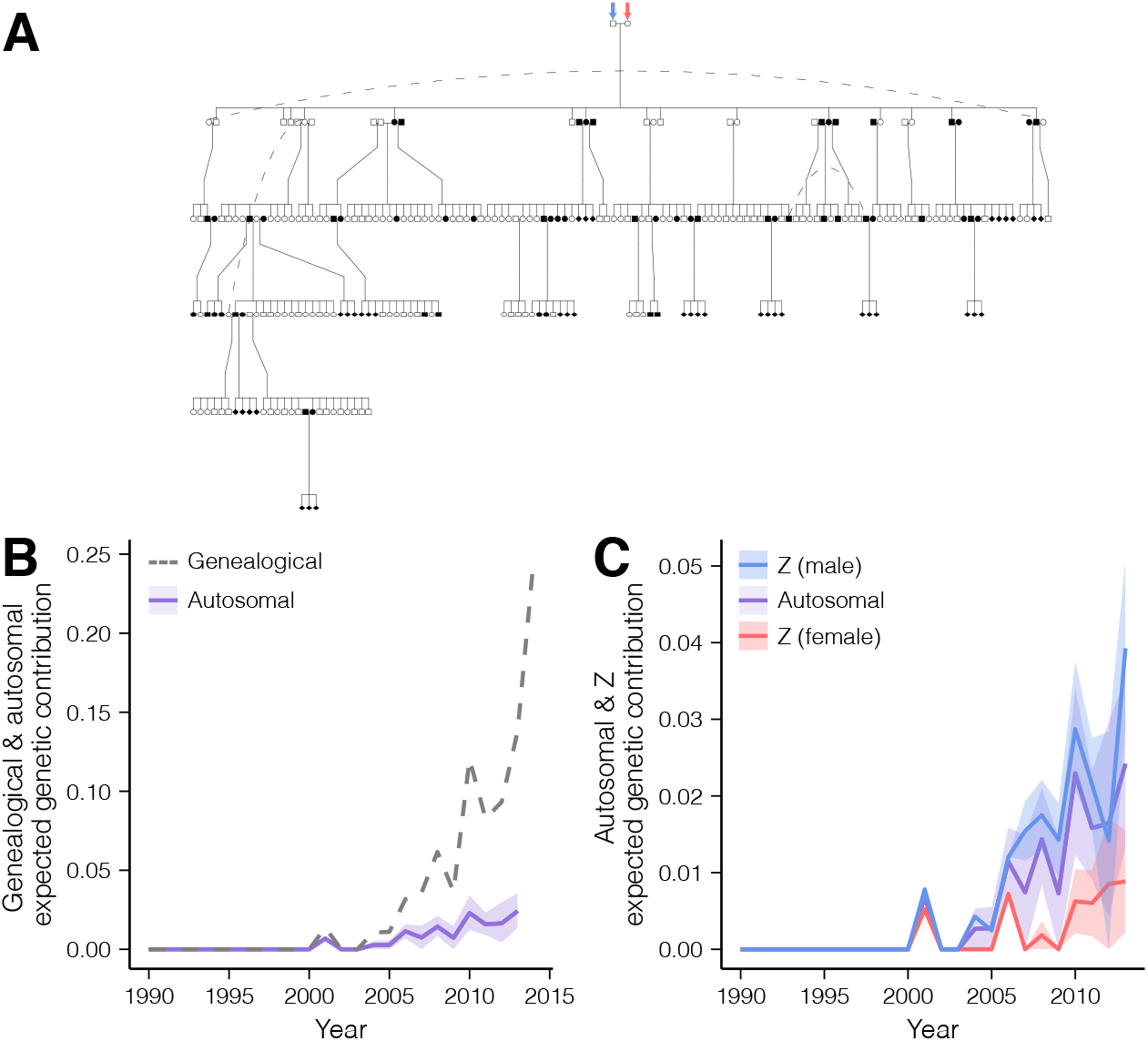
(A) Pedigree of descendants and (B, C) genealogical and genetic contributions over time for a male-female pair that first bred in 2001 with total lifetime reproductive success of 15. In A, dashed lines connect individuals who appear more than once in the pedigree. Solid symbols represent individuals still alive at end of study. In B, dashed lines indicate the proportion of nestlings in each cohort who are genealogical descendants of the pair. Solid lines indicate the mean expected genetic contribution at a neutral autosomal or Z chromosome locus for each year, and pale shading indicates the 95% confidence interval. Expected autosomal genetic contributions are shown in purple, and expected Z genetic contributions of the male partner are shown in blue and of the female in red.

Moving to the population level, we compared the genealogical and mean expected genetic contributions to our study population in 2013 of a set of 926 individuals who bred in our study population between 1990 and 2013. In this set of breeders, 317 males and 340 females have 0 descendants by 2013. For breeders with at least one descendant in 2013, genealogical contributions to the study population in 2013 ranged from 0.0034 to 0.31 (mean = 0.053) for males and from 0.0034 to 0.31 (mean = 0.054) for females. The similar range of genealogical contributions between males and females was unsurprising given this species is monogamous and the mortality rate is the same between the sexes (57, 75). Expected genetic contributions for an autosomal locus ranged from 0.00042 to 0.024 (mean = 0.0049) for both males and females. Expected genetic contributions for a Z chromosome locus ranged from 0 to 0.039 (mean = 0.0066) for males and from 0 to 0.021 (mean = 0.0037) for females. Differences in autosomal and Z expected genetic contributions of male and female partners depend on the number of mates an individual had throughout their lifetime and the sex ratio of their descendants (Supplemental Results, Fig. S4).

We found that for an autosomal locus, the relationship between genealogical and expected genetic contributions did not significantly differ for males versus females (Fig 2A; genealogical contribution: *p <* 2.2e-16; sex: *p* = 0.82; genealogical*sex: *p* = 0.16; Table S2). For a Z chromosome locus, on the other hand, the relationship between genealogical and expected genetic contributions for males versus females was strikingly different (Fig 2B; genealogical contribution: *p <* 2.2e-16; sex: *p* = 0.26; genealogical*sex : *p <* 2.2e-16). The relationship between genealogical and expected genetic contributions for females (genealogical*sex: *β* = 0.043, Standard Error (SE) = 0.004) was 48% lower than that for males (genealogical: *β* = 0.083, SE = 0.003; Table S2). For both autosomal and Z chromosome loci, expected genetic contributions were much lower than genealogical contributions, consistent with theoretical predictions (the gray dotted lines in Figs 2A and 2B show a 1:1 relationship).

**Figure 2.**
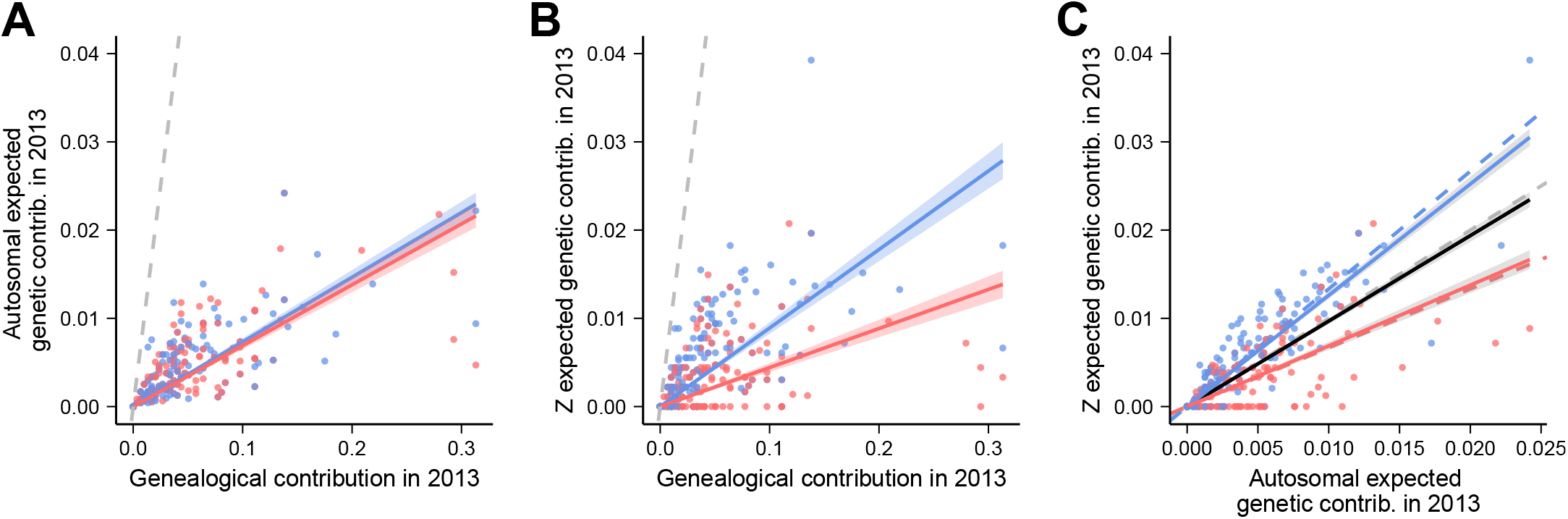
Genealogical and expected genetic contributions to the population in 2013 at (A) a neutral autosomal and (B) a neutral Z chromosome locus for all male breeders (*n* = 351, blue) and female breeders (*n* = 378, red) born before 2002 who bred in the population between 1990 and 2013. The dashed lines indicate a one-to-one relationship. (C) Expected genetic contributions to the population in 2013 at a neutral autosomal locus versus expected genetic contributions to the population in 2013 at a neutral Z chromosome locus for the breeders shown in parts A and B. The black line shows the Z/autosome relationship without regard for sex, the blue line shows the Z/autosome relationship for males, and the red line shows the Z/autosome relationship for females. Solid lines show estimates from linear models for each group, and dashed lines show theoretical expectations. Shading shows the standard error of the linear models.

We directly compared the expected genetic contributions of an autosomal locus and a Z locus. Assuming an equal sex ratio, the ratio of male Z/autosome expected genetic contributions is expected to be 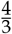 while female Z/autosome expected genetic contributions is expected to be 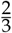 because males carry 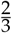 of the Z chromosomes and 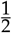 of the autosomes in a population. When sex is not considered, average expected genetic contributions to Z and autosomal loci should be equal because we defined expected genetic contribution as a proportion of the total number of Z chromosomes or autosomes respectively. As anticipated, the ratio of Z/autosome expected genetic contributions of individuals without regard to sex was close to 1 (slope = 0.97, SE = 0.018). When sex was added to the model (which explained 17% more variance than the model without sex), the Z/autosome ratio of expected genetic contributions was significantly affected by sex (*p <* 2.2e-16); the Z/autosome estimate for females (0.6747, SE = 0.034) was lower than the Z/autosome estimate for males (1.25, SE = 0.024) by 54%. The Z/autosome ratio of expected genetic contributions for females matched our expectations (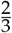), while the Z/autosome ratio for males was slightly less than the expected 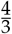 (Fig. 2C, Table S3).

### Expected genetic contributions of immigrants

Florida Scrub-Jays have female-biased dispersal, and previous work showed high levels of gene flow into our study population over time (16, 61). Between 1990 and 2012, the number of immigrant breeders appearing in Archbold each year was generally small (6-40/year; Fig. 3A). The immigrant breeder sex ratio was female-biased in 16 of 22 years (mean proportion of male immigrants across years = 0.39). Immigration decreased significantly over time (-0.41 *±* 0.06 immigrants/year, *R*^2^ = 0.34, *p*_*immigrant*_ = 2.61e-10; Table S4).

**Figure 3.**
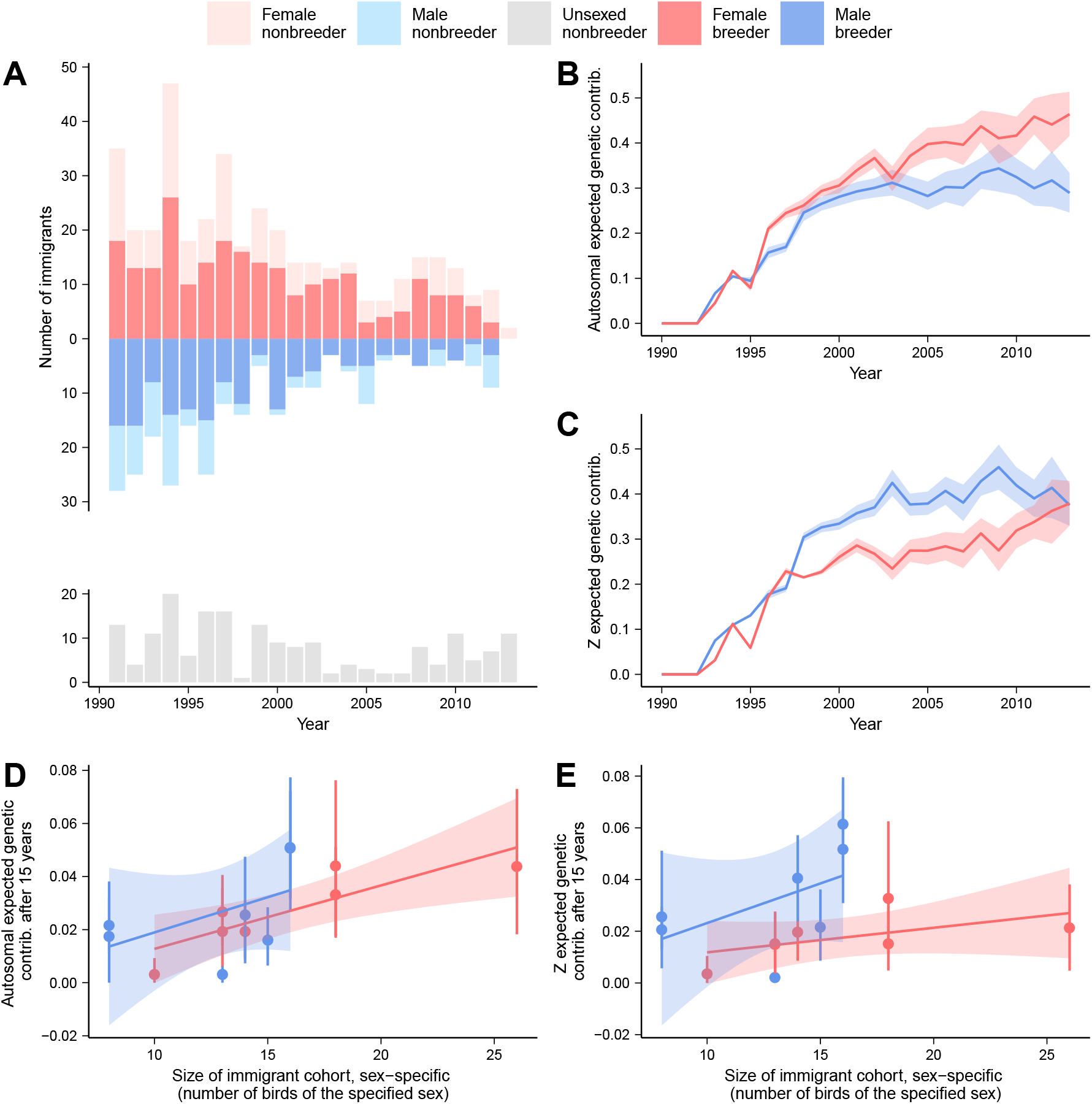
(A) The number of female immigrants (shown in light/dark red for nonbreeders/breeders), male immigrants (shown in light/dark blue for nonbreeders/breeders), and immigrants of unknown sex (all nonbreeders, shown in gray) arriving in the Archbold population each year between 1990 and 2013. (B, C) The expected genetic contribution of male and female immigrants appearing in the population after 1990 at (B) a neutral autosomal locus and (C) a neutral Z chromosome locus. Shading shows the 95% confidence interval. (D, E) The expected genetic contribution of immigrant cohorts between 1991 and 1997 to the nestling cohort 15 years later at (D) a neutral autosomal locus and (E) a neutral Z chromosome locus. Shading shows the standard error of the linear models and error bars indicate the 95% confidence interval.

To investigate the role of sex-biased migration, we calculated the expected genetic contributions of male and female immigrants who entered the population between 1990 and 2013. Female immigrants had significantly higher expected genetic contributions at autosomal loci than male immigrants by 2013 (Fig. 3B; significance based on non-overlapping 95% confidence intervals). However, despite female-biased immigration rates, the expected genetic contribution of male immigrants at Z-linked loci was significantly higher than that of female immigrants until 2011 (Fig. 3C). Consistent with previous work (16), 75% of the autosomal alleles in the 2013 birth cohort were contributed by immigrants arriving since 1990, 62% of which was driven by female immigrants (Table S5). On the Z chromosome, the total expected genetic contribution of male and female immigrants was also 75% in 2013 (Fig. 3C), with each sex contributing half (50%) of the incoming alleles. When we normalized expected Z genetic contributions by expected autosomal genetic contributions (thus correcting for sex-biased immigration), the Z/autosome ratio approached 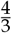 for male immigrants and 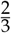 for female immigrants, as expected (Fig. S5, Table S5).

While all immigrants could make sustained genetic contributions to a population over time, gene flow could alternatively be driven by a few immigrant individuals with large contributions, in which case immigrant cohort size would not be a good predictor of gene flow. To this end, we assessed the relationship between immigrant cohort size and total expected genetic contribution of that immigrant cohort to individuals born 15 years later. We found that immigrant cohort size was a good predictor of autosomal (Fig 3D) and Z-linked (Fig 3E) expected genetic contributions. At autosomal loci, we found that the expected genetic contribution of a given immigrant cohort significantly increased with increasing cohort size, but there was no significant effect of sex on the rate of increase (*β* = 0.0025 *±* 0.0008, *R*^2^ = 0.35, *p*_*slope*_ = 0.012, *p*_*sex*_ = 0.34). Immigrant sex had an effect on Z-linked loci: immigrant cohort size was not significantly associated with expected Z genetic contribution after 15 years for female immigrants (*β* = 0.0016 *±* 0.00095, *p* = 0.1199) but was significant for male immigrants (*β* = 0.019 *±* 0.0084, *p* = 0.041; full model *R*^2^ = 0.25). Overall, we saw a similar relationship between immigrant cohort size and expected genetic contributions of male and female immigrant cohorts on the autosomes, but our results suggest male immigrant cohort size has a larger effect than female immigrant cohort size on expected genetic contributions on the Z.

### Signals of selection

We next evaluated signals of selection on 250 Z-linked loci by comparing the change in allele frequencies expected under the pedigree to the observed change (16). We identified loci at which the observed change in allele frequency between 1999 and 2013 was larger than expected from our simulations as potentially under directional selection. Across 1999-2013, we found no Z-linked SNPs that showed significantly larger shifts in allele frequency than would be expected from gene flow and drift alone at a FDR of 0.1 (1 SNP is significant at a less stringent FDR cutoff of 0.25; Fig. S6, Table S6). We also compared pairs of consecutive years (*e*.*g*., 1999 and 2000, 2000 and 2001, etc.) and found a total of 6 Z-linked SNPs—2 in 1999-2000, 3 in 2000-2001, and 1 in 2001-2002—that showed significantly more allele frequency change than would be expected from gene flow and drift alone from one year to the next at a FDR of 0.1 (Fig. S7, Table S6). All significant loci were Z-linked; none were in the pseudoautosomal region. We reassessed the 10,731 autosomal loci examined by (16) after correcting for multiple comparisons across both autosomal and Z loci and found results were largely unchanged. If we use the same FDR cutoff as (16) (0.25), the same 18 autosomal SNPs showed significantly larger shifts in allele frequency than would be expected from gene flow and drift alone across 1999-2013 (Table S7). For allele frequency shifts between adjacent years, 44 of the 47 previously-identified SNPs remained significant (3 SNPs in 2003-2004 were no longer significant; Table S8), and 4 new SNPs were identified (1 in 1999-2000 and 3 in 2000-2001). At a FDR < 0.1, we found no hits for autosomal SNPs across 1999-2013 and 9 hits across consecutive years: 7 SNPs in 2001-2002, 1 in 2010-2011, and 1 in 2012-2013. Overall, our results are consistent with previous findings that allele frequency changes in 1999-2013 are mostly consistent with a neutral model.

### Causes of allele frequency change

To quantify the relative roles of different evolutionary processes, we partitioned genome-wide allele frequency change between consecutive years among demographic groups (survivors, immigrants, nestlings/births; Fig. 4A), assuming survival/reproduction and gene flow are the only sources of allele frequency change. Building on the model of (16), we additionally assessed sex-specific demography and sex-specific inheritance by allowing differences between males and females in each demographic group (Fig. 4A) and creating separate models that use the transmission rules for the Z chromosome. Our models provide a good fit to our data: the predicted allele frequency change estimated from our model (*i*.*e*., the sum of each term in Eq. 3 and 4) is highly correlated with the observed allele frequency change for both the autosomes (Spearman’s *ρ* = 0.99, *p* < 2.2e-16) and the Z chromosome (Spearman’s *ρ* = 0.88, *p* < 2.2e-16; Fig. S8). Variance partition estimates obtained from the full dataset and an LD-pruned dataset of 4,200 SNPs are highly correlated (Spearman’s *ρ* = 0.99 for autosomes and 0.94 for the Z chromosome, *p* < 2.2e-16 for both; see Supplementary Material; Fig. S9).

**Figure 4.**
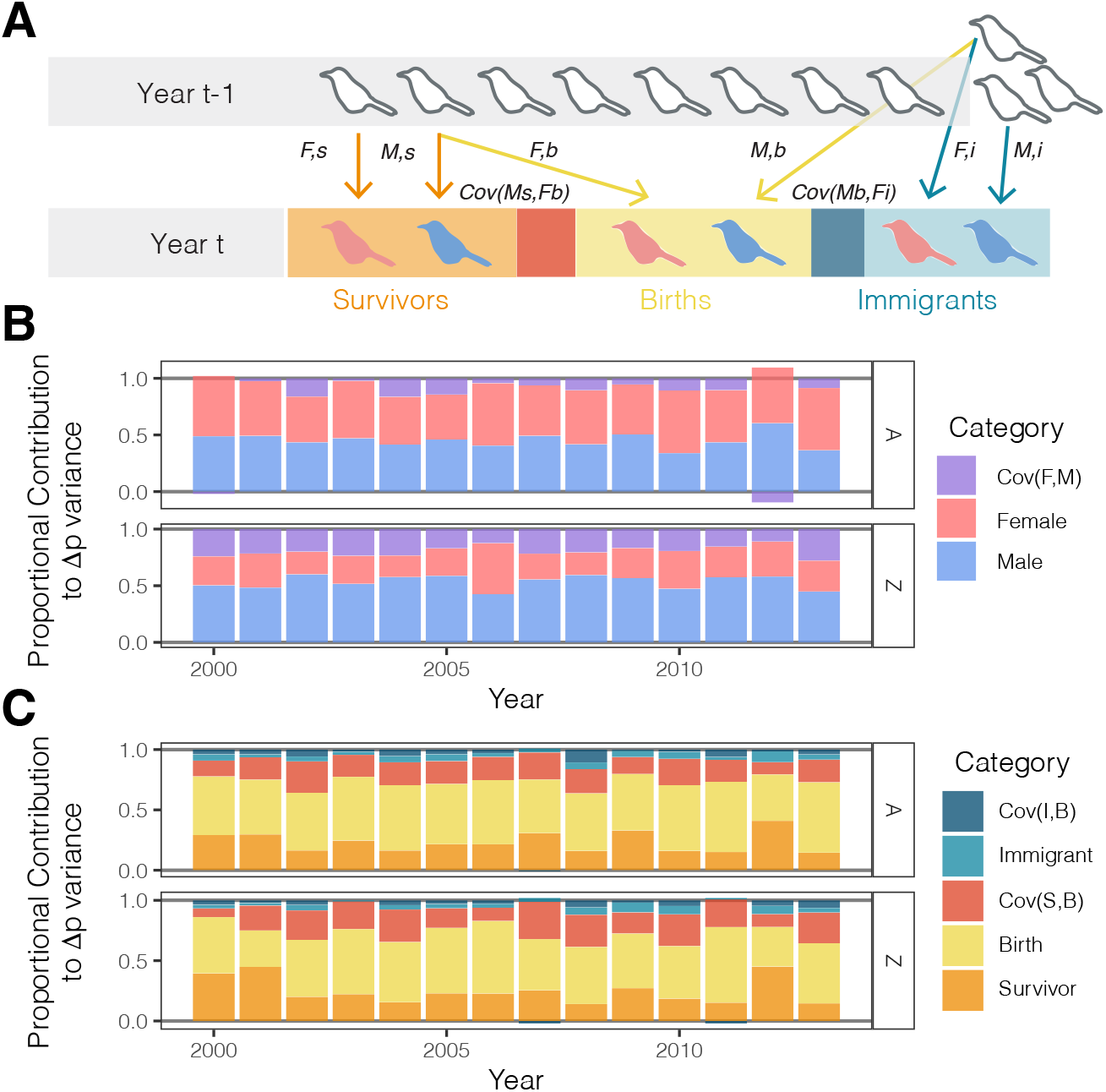
(A) Schematic of the allele frequency variance partitioning model. Arrows show contributions to allele frequency change through different demographic processes: survival, reproduction (birth), and immigration. *F,s* = female survivors; *M,s* = male survivors; *F,b* = female births; *M,b* = male births; *F,i* = female immigrants; and *M,i* = male immigrants. (B, C) Results of the model. (B) Allele frequency variance partitioning for males (blue), females (pink), and covariance between males and females (purple) for autosomal loci (top) and Z-linked loci (bottom). (C) Allele frequency variance partitioning for survivors (orange), births (yellow) and immigrants (teal), as well as covariance between survivors and births (Cov(S,B), red) and covariance between immigrants and births (Cov(I,B), blue) for autosomal loci (top) and Z-linked loci (bottom).

We first looked at the relative contributions of males, females, and male-female covariance to overall allele frequency change (Fig. 4B). For autosomes, the proportion of variance contributed by males and by females over years was generally similar, ranging from 0.34 to 0.60 for males and from 0.40 to 0.55 in females. Males had a similar contribution to variance in allele frequencies on the Z (0.42 to 0.60) while females contributed proportionally less to total allele frequency change on the Z (0.19 to 0.45) than on the autosomes. Note that the covariance between females and males is negative for autosomal loci in 2012. We also looked at the relative contributions of survivors, immigrants, births, and covariances between groups to overall allele frequency change (Fig. 4C). At autosomal loci, our findings were similar to the analysis of (16), with survivors, births, and survivor-birth covariance consistently contributing between 84 and 98% of the variance. Similarly, between 88 and 99% of the variance in allele frequencies on the Z chromosome was due to variation in survival, reproduction or survivor-birth covariance.

Next, we investigated the contributions of each sexed demographic group (Fig. 5). At autosomal loci, there was significant overlap in our confidence intervals between male and female survivors, suggesting both contribute similarly to allele frequency change from year to year. At Z-linked loci, male survivors tended to contribute more to variance in allele frequencies than females, but confidence intervals overlapped. We calculated the ratio between contributions to Z and autosomal allele frequency change and found that the Z/autosome ratio for female survivors approximated the expected 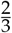, while the Z/autosome ratio for male survivors was more similar to 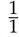 (Fig. S10). The covariance between male and female survivors declined across years on the Z, although it was positive in the first few years of the study period and confidence intervals overlapped zero in all years. Covariance between male and female survivors was consistently negative at autosomal loci, though confidence intervals again overlapped zero in all years (Fig. 5), and we showed that this observed negative covariance is a mathematical artifact (see Supplementary Material; Fig. S11).

**Figure 5.**
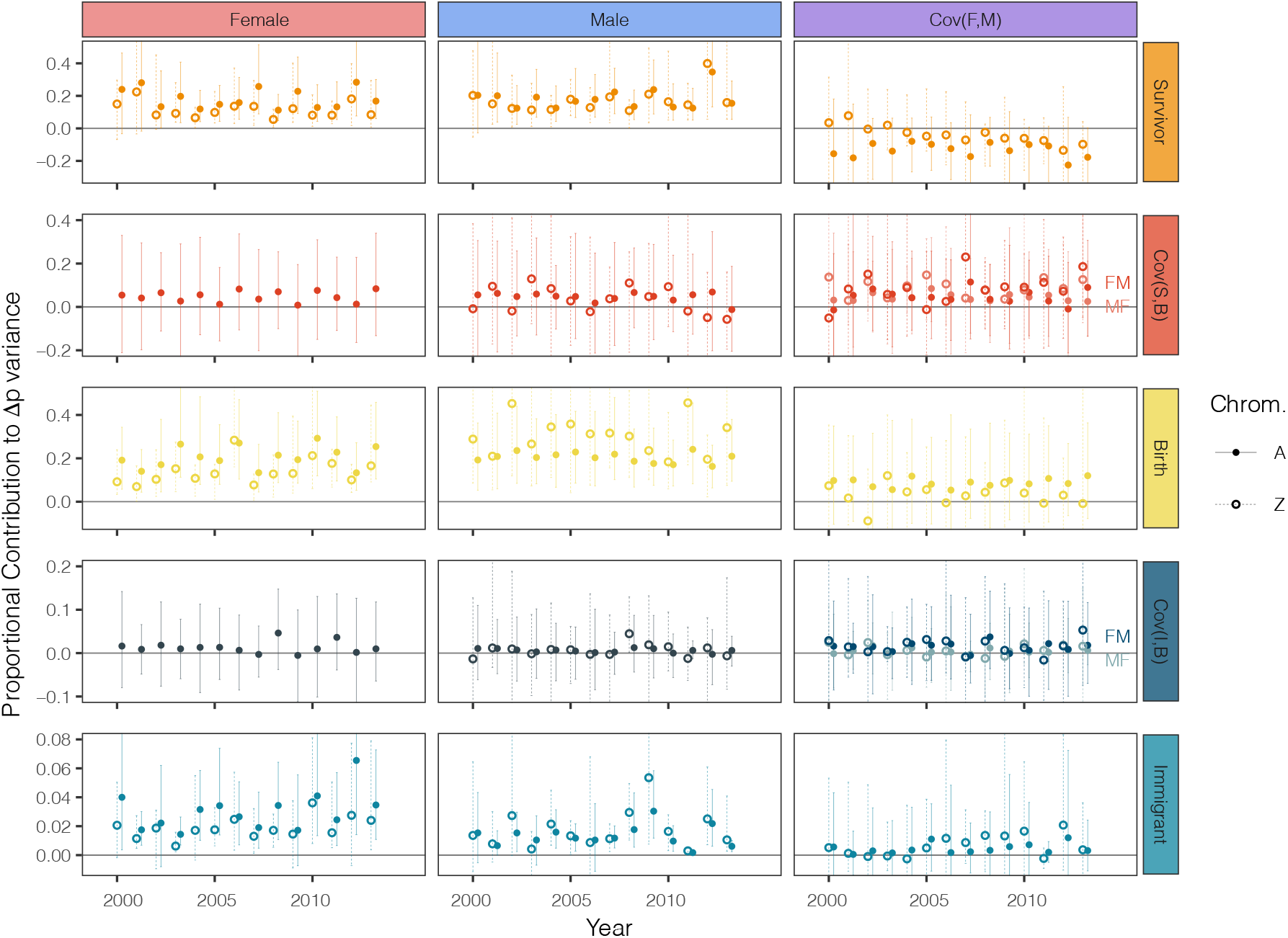
Contributions to the variance in allele frequency change across years of three demographic processes among the sexes. Solid circles and solid lines show the estimates for autosomal loci; open circles and dotted lines show the estimates for Z-linked loci. Vertical bars show 95% confidence intervals from bootstrapping. Cov(S,B) indicates covariance between survivors and births, and Cov(I,B) indicates covariance between immigrants and births. In the plot showing the covariance between survivors and births of different sexes, the lines labeled “MF” indicate cov(male survivors, female births) and the lines labeled “FM” indicate cov(female survivors, male births); likewise for the covariance between immigrants and births of different sexes. Note that the y-axis range varies among rows.

Confidence intervals for the variance in allele frequency change associated with male and female births overlapped for autosomal loci. On the Z, male births tended to play a larger role than female births, but confidence intervals overlapped here as well. For female births, the ratio between contributions to Z and autosomal allele frequency change was around 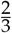, while for male births the ratio was around 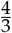 (Fig. S10). We further separated the variance associated with births into contributions of Mendelian segregation in heterozygous individuals and of variation in family size (Fig. S12). The ratio between contributions to Z and autosomal allele frequency change was approximately 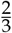 for both female and male Mendelian noise (Fig. S13). For female family size, the Z/autosome ratio was 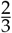, while for male family size the ratio was slightly above 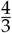. The covariance between male and female births was not significantly different from zero for both autosomes and the Z.

We found that, per year, immigration played a small role in allele frequency change (-0.0026 to 0.056), about an order of magnitude less than births (-0.088 to 0.50) or survival (-0.23 to 0.40). At autosomal loci, female immigrants tended to contribute more than males, although the confidence intervals overlapped. On the Z, male immigrants contributed more than females. The Z/autosome ratio for male immigrants was well above 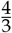, while the Z/autosome ratio for female immigrants was much lower than 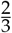 (Fig. S10).

## Discussion

The use of temporal samples to understand allele frequency change over time has been revolutionized by the advent of high-throughput genotyping technology. Here, we investigated the effects of sex-biased demography, sex-specific inheritance, and their interplay on expected genetic contributions and allele frequency change over short timescales using 11,000 SNPs and a multigenerational pedigree. Based on expected genetic contributions obtained by simulating the transmission of alleles down the pedigree, we found similar average contributions of the sexes on autosomes, but highly male-biased contributions to the Z chromosome. We also partitioned the variance in allele frequency change among consecutive years due to sex-specific survival/reproduction and gene flow. Consistent with previous work (16), we found that overall, differential birth and survival drives the majority of allele frequency change. While contributions to allele frequency change at autosomal loci are equally distributed between the sexes, males contribute more to allele frequency change at Z-linked loci than females. Together, our results offer unique insights into the impacts of sex-specific processes on evolution over ecologically relevant timescales.

We estimated an individual’s expected genetic contribution as the expected proportion of alleles in each birth cohort inherited identical-by-descent from the focal individual. After 10 generations, the expected genetic contributions of an individual should stabilize across years at the reproductive value for that individual (19, 76, 77). An individual’s reproductive value, a concept first introduced by Fisher (6) as a way to describe allele frequency change in age-structured populations, is obtained from an individual’s relative contribution to the future gene pool and is an accurate measure of fitness (19, 78). Reproductive values are traditionally estimated in population ecology as a weighted average of present and future reproduction by an individual at a given age (79). Although the standard concept of reproductive value described by Fisher assumes constant vital statistics (17), reproductive value can be measured for both individuals and classes (78), and we can define an individual’s reproductive value as its expected genetic contribution in distant generations conditional on its pedigree (19). Given that Florida Scrub-Jays have an estimated generation time of 5 years (80), we expect an individual Florida Scrub-Jay’s expected genetic contribution to converge to its reproductive value within 50 years (19). We found that the ratio of expected genetic contributions for autosomes and Z hover around 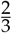 for females and 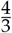 for males, suggesting that the expected genetic contributions of groups (*e*.*g*., all individuals of a given sex) can stabilize on much shorter timescales than the expected genetic contributions of individuals. Our work supports our growing theoretical understanding of differences in expected genetic contributions (81) and reproductive values on sex chromosomes (82, 83). Further understanding and quantifying the relationship between reproductive value on autosomes versus Z chromosomes is an important question for future work.

To quantify the contributions of different evolutionary processes and sexes to the variance in allele frequency change from year to year, we partitioned the variance in allele frequency change among sexes and demographic groups for autosomes and the Z chromosome. For the majority of the terms in our model, males contributed 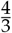 as much on the Z chromosome compared to autosomes, and females contributed 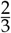 as much on the Z compared to autosomes. These ratios are strikingly similar to those expected from the differences in effective population size between autosomal and sex-linked regions of the genome (29). In fact, our results suggest that sex-biased survival or reproductive success have little effect on allele frequency change in the Florida Scrub-Jay; the only potentially sex-biased process is immigration. Indeed, the Florida Scrub-Jay is a simple case study from a population genetic standpoint. Florida Scrub-Jays are monogamous with a 50:50 breeder sex ratio and equal variance in reproductive success for males and females (56). Nonetheless, some terms in our allele frequency partitioning model departed from the 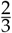 and 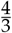 trend. First, when we separated allele frequency change due to births into the contributions of family size variation and Mendelian (random) assortment of chromosomes, the role of random assortment on the Z was 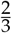 that of the autosomal term for both males and females (Fig. S13). This observation is to be expected as, for Z chromosomes, only one assortment event occurs per parent-offspring triad (during transmission from father to offspring), and thus all offspring regardless of sex have reduced change in allele frequencies on the Z chromosome compared to autosomes that is due to random of assortment of chromosomes. Second, the variance in allele frequency change contributed by the male survival term has a 1:1 Z/autosome ratio, which is lower than other ratios. There are several sources of variation that might affect this correlation, most notably the allele frequencies themselves, so we expect this correlation to be complex and are not presently able to pinpoint why this ratio is 1:1. Third, we observed no correlation between allele frequency change contributed by female immigrants on the Z and allele frequency change contributed by female immigrants on autosomes, in contrast to a strong correlation between Z and autosomal allele frequency variance in most categories. Future work could focus on the variation in Z/autosome ratios between demographic groups discovered here.

The Florida Scrub-Jay offers a useful test of the impact of sex-biased migration rates on short-term allele frequency dynamics: in our study population of jays, even though new immigrants (*i*.*e*., immigrants that just arrived that year) only make up a small proportion of the census population size in any given year, nearly a quarter of breeders in each year were once immigrants. Previous work on these jays (59) found higher identity-by-descent for the Z chromosome compared to the autosomes at short distances due to both a lower effective population size of the Z and female-biased dispersal. Here we showed that, due to sex-biased immigration, female immigrants have higher autosomal contributions than male immigrants. However, due to sex-biased ploidy and transmission of the Z, male immigrants had higher Z contributions than female immigrants, consistent with expectations. Our results reinforce the importance of maintaining/restoring connectivity among small populations otherwise vulnerable to allelic loss and inbreeding depression, and highlight how the sex ratio of translocated individuals may lead to different levels of genetic variation across the genome.

Our results also offer insights into the effect of sex-specific differences in variance in reproductive success. By following changes in allele frequency between years separately for males and females, we evaluated the effect of differences in life history between the sexes and their covariance on allele frequencies on the Z and autosomes. The general increase in the covariance between males and females on the Z compared to the autosomes is likely due to differences in transmission rules: because mothers do not transmit a Z chromosome to their daughters, daughters are more likely to share alleles with their brothers on the Z than on autosomes. However, by breaking down the contributions of covariances between males and females for survival and birth (reproduction) to allele frequency change, we may be able to evaluate whether sex-biases in reproductive success result in allele frequency change (84). In the Florida Scrub-Jay, we find the negative covariance in survival between the sexes is best explained by random changes in the adult sex ratio between years rather than by a systematic difference between the sexes. Future application of our method to systems with more complex mating systems, life histories, and demographies is likely to be highly informative of the propensity of sexual selection and sexual conflict to impact allele frequency changes on short timescales.

A more complete understanding of the effects of sex-biased processes on short-term evolutionary dynamics will be possible following the completion of a detailed linkage map and more dense genotyping, which would allow us to incorporate linkage into our analyses. Linkage disequilibrium can inflate the variance in individual realized genetic contributions (85). Linked selection influences Z to autosome diversity ratios because of different effective recombination rates on different chromosomes (86). The absence of recombination on the Z chromosome in the heterogametic sex (ZW in birds) causes the Z to have a lower effective recombination rate than the autosomes (all else being equal); these effects can be exacerbated by heterochiasmy. In the Florida Scrub-Jay, we have preliminary evidence that despite heterogeneity in sex differences in recombination rate across the genome, total map lengths of males and females are not significantly different. Here, we primarily focus on individual expected genetic contributions. Though our variance estimates for these expected values are likely a low estimate, we do not think linkage would significantly change the overall patterns we observe in expected genetic contributions of males and females. Future work that accounts for effective recombination rates and traces the inheritance of haplotypes down the pedigree to characterize actual, realized genetic contributions of individuals across the genome will provide a more detailed picture of how sex-biased demography and sex-biased transmission influence short-term evolutionary dynamics.

Linkage between neutral and selected loci affects our interpretation of signals of selection by distorting allele frequencies at neutral loci. Indeed, the hitchhiking of neutral alleles near sites under selection can make pinpointing the actual targets of selection challenging, until neutral and causal sites are separated by recombination (87). Thus, given our current method and SNP density, it is possible that some of the nine autosomal and six Z-linked SNPs that had larger than expected changes in allele frequency based on the pedigree are not the true targets of selection or (given the FDR of 0.1) are outlier loci whose frequency change is governed by chance. The lack of strong evidence for single SNPs under selection is unsurprising on our short timescale and in the relatively stable environment at Archbold Biological Station. Adaptation on short timescales often involves selection on polygenic traits or standing variation, which results in small allele frequency changes at many sites that are difficult to detect in individual SNP-based analyses.

Finally, linkage disequilibrium can inflate estimates of the variance in allele frequency change over time (88, 89). Recent work by (90, 91) capitalizes on the fact that linked selection increases the variance of neutral allele frequency change and creates temporal autocovariance in allele frequency change to quantify the genome-wide impact of polygenic linked selection. Their approach is similar to our variance partitioning of temporal allele frequency changes but cannot account for gene flow or overlapping generations (90, 91) (but see 92). By tracking allele frequency change in specific groups of individuals, we can estimate the contributions of gene flow and deal with overlapping generations. We found larger bootstrap confidence intervals on the Z chromosome compared to the autosomes (Wilcoxon rank sum test *W* = 21344, *p* = 0.008), though this increase is likely due to both lower recombination and fewer SNPs on the Z. While this study used only 250 Z-linked SNPs, we expect we effectively captured the history of the Z chromosome in the Florida Scrub-Jay because the average distance between our SNPs is smaller than the average breakdown in linkage disequilibrium. Allele frequency variance partitioning based on our full dataset of 11,000 SNPs and an LD-pruned dataset of 4,200 SNPs gave very similar results (Spearman’s *ρ* = 0.99 for autosomes and 0.94 for the Z chromosome, *p* < 2.2e-16 for both), suggesting that our results are fairly robust. However, future work using denser genotype data and incorporating information on recombination rate variation across the genome would provide more accurate estimates of contributions to the variance in allele frequency change.

Overall, we showed that both sex-biased dispersal and sex-biased transmission had a strong effect on Z chromosome dynamics in a population of Florida Scrub-Jays. We found that proportional contributions of males and females to Z chromosomes compared to autosomes follow a straightforward 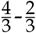 pattern in nearly every case, even on a relatively short evolutionary timescale. Our Beadchip dataset only captures common, variant sites; ongoing whole genome resequencing efforts will allow us to look at invariant sites and therefore tackle estimates of genetic diversity and test for a 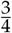 Z/autosome diversity pattern.

## Acknowledgments

Reed Bowman (1958-2023; Avian Ecology Program, Archbold Biological Station, Venus, FL 33960) was a co-author on this project. Faye Romero uploaded the genome assembly to NCBI. We would like to thank Graham Coop for his help with developing the allele frequency partitioning model. We would also like to thank the Chen and Coop labs, Nick Barton, two anonymous reviewers, and Ben Peter for feedback on the manuscript. Thank you to the many students and staff at Archbold Biological Station who collected field data.

