## Appendices for "Allele frequency dynamics under sex-biased demography and sex-specific inheritance in a pedigreed jay population"

### Appendix A. Derivations for allele frequency change variance partitioning models

#### Autosomal loci

##### Allele frequency variance model

We construct a model of allele frequency change over time that accounts for the overlapping generations and fluctuating population sizes characteristic of the Archbold population of Florida Scrub-Jays. Our model is based on the model of [1], but here we divide each demographic group (survivors, immigrants, births) into separate components for males and females. We model autosomal and Z-linked loci separately. For autosomal loci, let $N_{M,s}$ be the number of males who survived from year $t-1$ to year $t$. Let $N_{M,i}$ be the number of males who immigrated into the population in year $t$. Let $N_{M,b}$ be the number of males born in the population in year $t$. Let $N_{F,s}$ be the number of females who survived from year $t-1$ to year $t$. Let $N_{F,i}$ be the number of females who immigrated into the population in year $t$. Let $N_{F,b}$ be the number of females born in the population in year $t$. All individuals in the population in a given year fit into one of these six categories (male or female survivors, immigrants, or births) so $N_t = N_{F,s}+N_{F,i}+N_{F,b}+N_{M,s}+N_{M,i}+N_{M,b}$. If we represent the true allele counts for each sex ($s$) in each demographic group ($j$) as $P_{s,j}$, then the true allele frequencies can be represented as $p_{s,j} = \frac{P_{s,j}}{2N_{2,j}}$. The change in allele frequencies from year to year for autosomal SNPs is then:

$$
\begin{aligned}
\Delta p = p_t - p_{t-1} &= \frac{P_t}{2N_t} - \frac{P_{t-1}}{2N_{t-1}} = \frac{P_F + P_M}{2N_t} - \frac{P_{t-1}}{2N_{t-1}} \\
&= \frac{P_{F,s}+P_{F,i}+P_{F,b}+P_{M,s}+P_{M,i}+P_{M,b}}{2N_t} - \frac{P_{t-1}}{2N_{t-1}} \\
&= \frac{P_{F,s}+P_{F,i}+P_{F,b}+P_{M,s}+P_{M,i}+P_{M,b}}{2N_t}\left(\frac{N_{t-1}}{N_{t-1}}\right) \\
&\quad - \frac{P_{t-1}}{2N_{t-1}}\left(\frac{N_{F,s}+N_{F,i}+N_{F,b}+N_{M,s}+N_{M,i}+N_{M,b}}{N_t}\right) \\
&= \frac{P_{F,s}N_{t-1} - P_{t-1}N_{F,s}}{2N_tN_{t-1}} + \frac{P_{F,i}N_{t-1} - P_{t-1}N_{F,i}}{2N_tN_{t-1}} + \frac{P_{F,b}N_{t-1} - P_{t-1}N_{F,b}}{2N_tN_{t-1}} \\
&\quad + \frac{P_{M,s}N_{t-1} - P_{t-1}N_{M,s}}{2N_tN_{t-1}} + \frac{P_{M,i}N_{t-1} - P_{t-1}N_{M,i}}{2N_tN_{t-1}} + \frac{P_{M,b}N_{t-1} - P_{t-1}N_{M,b}}{2N_tN_{t-1}} \\
&= \left(\frac{N_{F,s}}{N_{F,s}}\right)\frac{P_{F,s}N_{t-1} - P_{t-1}N_{F,s}}{2N_tN_{t-1}} + \left(\frac{N_{F,i}}{N_{F,i}}\right)\frac{P_{F,i}N_{t-1} - P_{t-1}N_{F,i}}{2N_tN_{t-1}} + \left(\frac{N_{F,b}}{N_{F,b}}\right)\frac{P_{F,b}N_{t-1} - P_{t-1}N_{F,b}}{2N_tN_{t-1}} \\
&\quad + \left(\frac{N_{M,s}}{N_{M,s}}\right)\frac{P_{M,s}N_{t-1} - P_{t-1}N_{M,s}}{2N_tN_{t-1}} + \left(\frac{N_{M,i}}{N_{M,i}}\right)\frac{P_{M,i}N_{t-1} - P_{t-1}N_{M,i}}{2N_tN_{t-1}} + \left(\frac{N_{M,b}}{N_{M,b}}\right)\frac{P_{M,b}N_{t-1} - P_{t-1}N_{M,b}}{2N_tN_{t-1}} \\
&= \left(\frac{P_{F,s}N_{F,s}}{2N_tN_{F,s}} - \frac{P_{t-1}N_{F,s}}{2N_tN_{t-1}}\right) + \left(\frac{P_{F,i}N_{F,i}}{2N_tN_{F,i}} - \frac{P_{t-1}N_{F,i}}{2N_tN_{t-1}}\right) + \left(\frac{P_{F,b}N_{F,b}}{2N_tN_{F,b}} - \frac{P_{t-1}N_{F,b}}{2N_tN_{t-1}}\right) \\
&\quad + \left(\frac{P_{M,s}N_{M,s}}{2N_tN_{M,s}} - \frac{P_{t-1}N_{M,s}}{2N_tN_{t-1}}\right) + \left(\frac{P_{M,i}N_{M,i}}{2N_tN_{M,i}} - \frac{P_{t-1}N_{M,i}}{2N_tN_{t-1}}\right) + \left(\frac{P_{M,b}N_{M,b}}{2N_tN_{M,b}} - \frac{P_{t-1}N_{M,b}}{2N_tN_{t-1}}\right) \\
&= \frac{N_{F,s}}{N_t}\left(\frac{P_{F,s}}{2N_{F,s}} - \frac{P_{t-1}}{2N_{t-1}}\right) + \frac{N_{F,i}}{N_t}\left(\frac{P_{F,i}}{2N_{F,i}} - \frac{P_{t-1}}{2N_{t-1}}\right) + \frac{N_{F,b}}{N_t}\left(\frac{P_{F,b}}{2N_{F,b}} - \frac{P_{t-1}}{2N_{t-1}}\right) \\
&\quad + \frac{N_{M,s}}{N_t}\left(\frac{P_{M,s}}{2N_{M,s}} - \frac{P_{t-1}}{2N_{t-1}}\right) + \frac{N_{M,i}}{N_t}\left(\frac{P_{M,i}}{2N_{M,i}} - \frac{P_{t-1}}{2N_{t-1}}\right) + \frac{N_{M,b}}{N_t}\left(\frac{P_{M,b}}{2N_{M,b}} - \frac{P_{t-1}}{2N_{t-1}}\right) \\
&= \frac{N_{F,s}}{N_t}(p_{F,s} - p_{t-1}) + \frac{N_{F,i}}{N_t}(p_{F,i} - p_{t-1}) + \frac{N_{F,b}}{N_t}(p_{F,b} - p_{t-1}) \\
&\quad + \frac{N_{M,s}}{N_t}(p_{M,s} - p_{t-1}) + \frac{N_{M,i}}{N_t}(p_{M,i} - p_{t-1}) + \frac{N_{M,b}}{N_t}(p_{M,b} - p_{t-1}) \qquad (1)
\end{aligned}
$$

The variance in allele frequency change over time is then:

$$
\begin{aligned}
\mathrm{Var}(\Delta p) = \mathrm{Var}\Bigg[ & \frac{N_{F,s}}{N_t}(p_{F,s} - p_{t-1}) + \frac{N_{F,i}}{N_t}(p_{F,i} - p_{t-1}) + \frac{N_{F,b}}{N_t}(p_{F,b} - p_{t-1}) \\
& + \frac{N_{M,s}}{N_t}(p_{M,s} - p_{t-1}) + \frac{N_{M,i}}{N_t}(p_{M,i} - p_{t-1}) + \frac{N_{M,b}}{N_t}(p_{M,b} - p_{t-1})\Bigg] \\
= & \left(\frac{N_{F,s}}{N_t}\right)^2 \mathrm{Var}(p_{F,s} - p_{t-1}) + \left(\frac{N_{F,i}}{N_t}\right)^2 \mathrm{Var}(p_{F,i} - p_{t-1}) + \left(\frac{N_{F,b}}{N_t}\right)^2 \mathrm{Var}(p_{F,b} - p_{t-1}) \\
& + \left(\frac{N_{M,s}}{N_t}\right)^2 \mathrm{Var}(p_{M,s} - p_{t-1}) + \left(\frac{N_{M,i}}{N_t}\right)^2 \mathrm{Var}(p_{M,i} - p_{t-1}) + \left(\frac{N_{M,b}}{N_t}\right)^2 \mathrm{Var}(p_{M,b} - p_{t-1}) \\
& + 2\frac{N_{F,s}N_{F,b}}{N_t^2}\mathrm{Cov}(p_{F,s} - p_{t-1}, p_{F,b} - p_{t-1}) + 2\frac{N_{F,s}N_{M,s}}{N_t^2}\mathrm{Cov}(p_{F,s} - p_{t-1}, p_{M,s} - p_{t-1}) \\
& + 2\frac{N_{F,s}N_{M,b}}{N_t^2}\mathrm{Cov}(p_{F,s} - p_{t-1}, p_{M,b} - p_{t-1}) + 2\frac{N_{F,i}N_{F,b}}{N_t^2}\mathrm{Cov}(p_{F,i} - p_{t-1}, p_{F,b} - p_{t-1}) \\
& + 2\frac{N_{F,i}N_{M,i}}{N_t^2}\mathrm{Cov}(p_{F,i} - p_{t-1}, p_{M,i} - p_{t-1}) + 2\frac{N_{F,i}N_{M,b}}{N_t^2}\mathrm{Cov}(p_{F,i} - p_{t-1}, p_{M,b} - p_{t-1}) \\
& + 2\frac{N_{F,b}N_{M,s}}{N_t^2}\mathrm{Cov}(p_{F,b} - p_{t-1}, p_{M,s} - p_{t-1}) + 2\frac{N_{F,b}N_{M,i}}{N_t^2}\mathrm{Cov}(p_{F,b} - p_{t-1}, p_{M,i} - p_{t-1}) \\
& + 2\frac{N_{F,b}N_{M,b}}{N_t^2}\mathrm{Cov}(p_{F,b} - p_{t-1}, p_{M,b} - p_{t-1}) + 2\frac{N_{M,s}N_{M,b}}{N_t^2}\mathrm{Cov}(p_{M,s} - p_{t-1}, p_{M,b} - p_{t-1}) \\
& + 2\frac{N_{M,i}N_{M,b}}{N_t^2}\mathrm{Cov}(p_{M,i} - p_{t-1}, p_{M,b} - p_{t-1}) \qquad (2)
\end{aligned}
$$

Our model partitions the variance in allele frequency change into contributions from female survivors, female immigrants, female births, male survivors, male immigrants, male births, and their covariances. We assume immigrants in a given year are unrelated to survivors and therefore set covariances between survivors and immigrants to 0.

#### Estimation of variance due to Mendelian segregation

The variance in allele frequency change due to births derives from the effects of both variation in family sizes and Mendelian segregation of alleles from heterozygous parents. Here, we show how the variance in allele frequency change due to these two sources of variation can be partitioned.

In year $t$, the allele frequency of the birth cohort at a given autosomal locus is the sum of the allele frequency ($g_{s,k}$) of each individual nestling ($k$). We model this separately for each sex, $s$, as follows:

$$p_{s,b} = \sum_{k=1}^{N_{s,b}} g_{s,k} \qquad (3)$$

where $g_{s,k}$ has a value of 0, 0.5, or 1 for individuals with genotype 00, 01, or 11 respectively. We can express $g_{s,k}$ as:

$$g_{s,k} = \frac{1}{2}(g_{k,dad} + g_{k,mom}) + \frac{1}{2}(\delta_{k,dad} + \delta_{k,mom}) \qquad (4)$$

where $g_{k,dad}$ and $g_{k,mom}$ are the allele frequencies of the father and mother of individual $k$. Each is multiplied by $\frac{1}{2}$ as each parent contributes one allele to the offspring. $\delta_{k,dad}$ and $\delta_{k,mom}$ are the difference in allele frequency between individual $k$ and its parents due to Mendelian segregation.

The allele frequency of the birth cohort of each sex in a given year is then:

$$p_{s,b} = \sum_{k=1}^{N_{s,b}} \left[ \frac{1}{2}(g_{k,dad} + g_{k,mom}) + \frac{1}{2}(\delta_{k,dad} + \delta_{k,mom}) \right]$$
$$= \frac{1}{2}(p_{s,dad} + p_{s,mom}) + \frac{1}{2} \sum_{k=1}^{N_{s,b}} (\delta_{k,dad} + \delta_{k,mom}) \tag{5}$$

where $p_{s,dad}$ is the allele frequency of the fathers of nestlings of sex $s$ born that year, weighted by the number of offspring of that sex each father produced, and $p_{s,mom}$ is the allele frequency of the mothers of sex $s$ nestlings born that year, similarly weighted by the number of offspring of that sex each mother produced. The allele frequency of the birth cohort depends on both family size effects ($p_{s,b,fam}$) and the Mendelian segregation of alleles in heterozygotes ($p_{s,b,mend}$). Let us denote the first term as $p_{s,b,fam}$ and the second term as $p_{s,b,mend}$. We can now express $\text{Var}(p_{s,b} - p_{t-1})$ in terms of the variance due to family size and Mendelian noise:

$$\text{Var}(p_{s,b} - p_{t-1}) = \text{Var}(p_{s,b,fam} + p_{s,b,mend} - p_{t-1})$$
$$= \text{Var}(p_{s,b,fam} - p_{t-1}) + \text{Var}(p_{s,b,mend}) \tag{6}$$

To estimate the variance due to Mendelian noise, we solve for $p_{s,b,mend}$ :

$$\text{Var}(p_{s,b,mend}) = \text{Var}\left[ p_{s,b} - \frac{1}{2}(p_{s,dad} + p_{s,mom}) \right] \tag{7}$$

**Sampling error**

We know the number of individuals in each category in each year ($N_j$), but not all of these individuals are genotyped. Thus, the true value of $p_j$ is unknown. We correct for sampling error as follows. Let $n_j$ be the number of genotyped individuals in a given category in a given year and $x_j$ be the observed allele frequency. The difference between the observed and true allele frequencies — the error due to sampling — may then be represented for both males and females by $\varepsilon_j$ such that $p_j = x_j - \varepsilon_j$. The true allele frequencies $p_j$ in our model may now be broken down into observed allele frequencies and sampling error. Using male survivors as an example:

$$p_{M,s} - p_{t-1} = (x_{M,s} - \varepsilon_{M,s}) - (x_{t-1} - \varepsilon_{t-1}) = (x_{M,s} - x_{t-1}) - (\varepsilon_{M,s} - \varepsilon_{t-1}) \tag{8}$$

and

$$\text{Var}(p_{M,s} - p_{t-1}) = \text{Var}(x_{M,s} - x_{t-1}) - \text{Var}(\varepsilon_{M,s} - \varepsilon_{t-1}) - 2\text{Cov}(p_{M,s} - p_{t-1}, \varepsilon_{M,s} - \varepsilon_{t-1}) \tag{9}$$

The other variance terms break down similarly. For the covariance terms, we will use the covariance between male survivors and male births as an example:

$$\text{Cov}\,(p_{M,s} - p_{t-1}, p_{M,b} - p_{t-1}) = \text{Cov}(x_{M,s} - \varepsilon_{M,s} - x_{t-1} + \varepsilon_{t-1}, x_{M,b} - \varepsilon_{M,b} - x_{t-1} + \varepsilon_{t-1})$$
$$= \text{Cov}(x_{M,s} - x_{t-1}, x_{M,b} - x_{t-1})$$
$$- \text{Cov}(x_{M,s} - x_{t-1}, \varepsilon_{M,b} - \varepsilon_{t-1})$$
$$- \text{Cov}(\varepsilon_{M,s} - \varepsilon_{t-1}, x_{M,b} - x_{t-1})$$
$$+ \text{Cov}(\varepsilon_{M,s} - \varepsilon_{t-1}, \varepsilon_{M,b} - \varepsilon_{t-1}) \tag{10}$$

The remaining eleven covariance terms break down similarly.

We also incorporate error due to sampling into our estimate for Mendelian noise and family size:

$$\begin{aligned}
\mathrm{Var}(p_{s,b,mend}) &= \mathrm{Var}\left[x_{s,b} - \varepsilon_{s,b} - \frac{1}{2}(x_{s,dad} - \varepsilon_{s,dad} + x_{s,mom} - \varepsilon_{s,mom})\right] \\
&= \mathrm{Var}\left[x_{s,b} - \frac{1}{2}\left(x_{s,dad} + x_{s,mom}\right)\right] - \mathrm{Var}\left[\varepsilon_{s,b} - \frac{1}{2}\left(\varepsilon_{s,dad} + \varepsilon_{s,mom}\right)\right] \qquad (11) \\
\mathrm{Var}(p_{s,b,fam} - p_{t-1}) &= \mathrm{Var}\left[\frac{1}{2}(p_{s,dad} + p_{s,mom}) - p_{t-1}\right] \\
&= \mathrm{Var}\left[\frac{1}{2}(x_{s,dad} - \varepsilon_{s,dad} + x_{s,mom} - \varepsilon_{s,mom}) - (x_{t-1} - \varepsilon_{t-1})\right] \\
&= \mathrm{Var}\left[\frac{1}{2}(x_{s,dad} + x_{s,mom}) - x_{t-1}\right] - \mathrm{Var}\left[\frac{1}{2}(\varepsilon_{s,dad} + \varepsilon_{s,mom}) - \varepsilon_{t-1}\right] \qquad (12)
\end{aligned}$$

Throughout, we assume that the covariance between the observed allele frequency and the error is 0.

### Z-linked loci

#### Allele frequency variance model

We use a similar approach to model allele frequency change for Z-linked loci, but redefine a few terms to account for the unique transmission rules of the Z chromosome. For males, let $N_{M,t}$ be the total number of males in the population in year $t$. Let $N_{M,s}$ be the number of males who survived from year $t-1$ to year $t$. Let $N_{M,i}$ be the number of males who immigrated into the population in year $t$. Let $N_{M,b}$ be the number of males born in the population in year $t$. All males in the population in a given year fit into one of these three categories (survivors, immigrants, or births) so $N_{M,t} = N_{M,s} + N_{M,i} + N_{M,b}$. If we represent the true allele counts for males in each category as $P_{M,j}$, then the true allele frequencies can be represented as $p_{M,j} = \frac{P_{M,j}}{2N_{M,j}}$ because each male has two Z chromosomes.

Likewise, for females, let $N_{F,t}$ be the total number of females in the population in year $t$; $N_{F,s}$ the number of females who survived from year $t-1$ to year $t$; $N_{F,i}$ the number of females who immigrated into the population in year $t$; and $N_{F,b}$ the number of females born in the population in year $t$. As with males, $N_{F,t} = N_{F,s} + N_{F,i} + N_{F,b}$. If we represent the true allele counts for females in each category as $P_{F,j}$, then the true allele frequencies can be represented as $p_{F,j} = \frac{P_{F,j}}{N_{F,j}}$ because each female has just one Z chromosome.

The change in allele frequencies from year to year for Z-linked SNPs is:

$$
\begin{aligned}
\Delta p &= p_t - p_{t-1} \\
&= \frac{P_t}{N_{F,t} + 2N_{M,t}} - \frac{P_{t-1}}{N_{F,t-1} + 2N_{M,t-1}} \\
&= \frac{P_{F,s} + P_{F,i} + P_{F,b} + P_{M,s} + P_{M,i} + P_{M,b}}{N_{F,t} + 2N_{M,t}} \\
&\quad - \frac{P_{t-1}}{N_{F,t-1} + 2N_{M,t-1}} \left( \frac{N_{F,s} + N_{F,i} + N_{F,b} + 2N_{M,s} + 2N_{M,i} + 2N_{M,b}}{N_{F,t} + 2N_{M,t}} \right) \\
&= \left[ \frac{P_{F,s}}{N_{F,t} + 2N_{M,t}} \left( \frac{N_{F,s}}{N_{F,s}} \right) + \frac{P_{F,i}}{N_{F,t} + 2N_{M,t}} \left( \frac{N_{F,i}}{N_{F,i}} \right) + \frac{P_{F,b}}{N_{F,t} + 2N_{M,t}} \left( \frac{N_{F,b}}{N_{F,b}} \right) \right. \\
&\quad \left. + \frac{P_{M,s}}{N_{F,t} + 2N_{M,t}} \left( \frac{2N_{M,s}}{2N_{M,s}} \right) + \frac{P_{M,i}}{N_{F,t} + 2N_{M,t}} \left( \frac{2N_{M,i}}{2N_{M,i}} \right) + \frac{P_{M,b}}{N_{F,t} + 2N_{M,t}} \left( \frac{2N_{M,b}}{2N_{M,b}} \right) \right] \\
&\quad - \left[ \frac{P_{t-1}N_{F,s}}{(N_{F,t} + 2N_{M,t})(N_{F,t-1} + 2N_{M,t-1})} + \frac{P_{t-1}N_{F,i}}{(N_{F,t} + 2N_{M,t})(N_{F,t-1} + 2N_{M,t-1})} \right. \\
&\quad + \frac{P_{t-1}N_{F,b}}{(N_{F,t} + 2N_{M,t})(N_{F,t-1} + 2N_{M,t-1})} + \frac{P_{t-1}2N_{M,s}}{(N_{F,t} + 2N_{M,t})(N_{F,t-1} + 2N_{M,t-1})} \\
&\quad \left. + \frac{P_{t-1}2N_{M,i}}{(N_{F,t} + 2N_{M,t})(N_{F,t-1} + 2N_{M,t-1})} + \frac{P_{t-1}2N_{M,b}}{(N_{F,t} + 2N_{M,t})(N_{F,t-1} + 2N_{M,t-1})} \right] \\
&= \frac{P_{F,s}N_{F,s}}{(N_{F,t} + 2N_{M,t})N_{F,s}} - \frac{P_{t-1}N_{F,s}}{(N_{F,t} + 2N_{M,t})(N_{F,t-1} + 2N_{M,t-1})} \\
&\quad + \frac{P_{F,i}N_{F,i}}{(N_{F,t} + 2N_{M,t})N_{F,i}} - \frac{P_{t-1}N_{F,i}}{(N_{F,t} + 2N_{M,t})(N_{F,t-1} + 2N_{M,t-1})} \\
&\quad + \frac{P_{F,b}N_{F,t}}{(N_{F,t} + 2N_{M,t})N_{F,t}} - \frac{P_{t-1}N_{F,b}}{(N_{F,t} + 2N_{M,t})(N_{F,t-1} + 2N_{M,t-1})} \\
&\quad + \frac{P_{M,s}2N_{M,s}}{(N_{F,t} + 2N_{M,t})2N_{M,s}} - \frac{P_{t-1}2N_{M,s}}{(N_{F,t} + 2N_{M,t})(N_{F,t-1} + 2N_{M,t-1})} \\
&\quad + \frac{P_{M,i}2N_{M,i}}{(N_{F,t} + 2N_{M,t})2N_{M,i}} - \frac{P_{t-1}2N_{M,i}}{(N_{F,t} + 2N_{M,t})(N_{F,t-1} + 2N_{M,t-1})} \\
&\quad + \frac{P_{M,b}2N_{M,b}}{(N_{F,t} + 2N_{M,t})2N_{M,b}} - \frac{P_{t-1}2N_{M,b}}{(N_{F,t} + 2N_{M,t})(N_{F,t-1} + 2N_{M,t-1})} \\
&= \frac{N_{F,s}}{N_{F,t} + 2N_{M,t}} \left( \frac{P_{F,s}}{N_{F,s}} - \frac{P_{t-1}}{N_{F,t-1} + 2N_{M,t-1}} \right) \\
&\quad + \frac{N_{F,i}}{N_{F,t} + 2N_{M,t}} \left( \frac{P_{F,i}}{N_{F,i}} - \frac{P_{t-1}}{N_{F,t-1} + 2N_{M,t-1}} \right) \\
&\quad + \frac{N_{F,b}}{N_{F,t} + 2N_{M,t}} \left( \frac{P_{F,b}}{N_{F,b}} - \frac{P_{t-1}}{N_{F,t-1} + 2N_{M,t-1}} \right) \\
&\quad + \frac{2N_{M,s}}{N_{F,t} + 2N_{M,t}} \left( \frac{P_{M,s}}{2N_{M,s}} - \frac{P_{t-1}}{N_{F,t-1} + 2N_{M,t-1}} \right) \\
&\quad + \frac{2N_{M,i}}{N_{F,t} + 2N_{M,t}} \left( \frac{P_{M,i}}{2N_{M,i}} - \frac{P_{t-1}}{N_{F,t-1} + 2N_{M,t-1}} \right) \\
&\quad + \frac{2N_{M,b}}{N_{F,t} + 2N_{M,t}} \left( \frac{P_{M,b}}{2N_{M,b}} - \frac{P_{t-1}}{N_{F,t-1} + 2N_{M,t-1}} \right) \\
&= \frac{N_{F,s}}{N_{F,t} + 2N_{M,t}} (p_{F,s} - p_{t-1}) + \frac{N_{F,i}}{N_{F,t} + 2N_{M,t}} (p_{F,i} - p_{t-1}) + \frac{N_{F,b}}{N_{F,t} + 2N_{M,t}} (p_{F,b} - p_{t-1}) \\
&\quad + \frac{2N_{M,s}}{N_{F,t} + 2N_{M,t}} (p_{M,s} - p_{t-1}) + \frac{2N_{M,i}}{N_{F,t} + 2N_{M,t}} (p_{M,i} - p_{t-1}) + \frac{2N_{M,b}}{N_{F,t} + 2N_{M,t}} (p_{M,b} - p_{t-1}) \qquad (13)
\end{aligned}
$$

The variance in allele frequency change over time is then:

$$\begin{aligned}
\mathrm{Var}(\Delta p) = \mathrm{Var}\Bigg[ & \frac{N_{F,s}}{N_{F,t}+2N_{M,t}}(p_{F,s}-p_{t-1}) + \frac{N_{F,i}}{N_{F,t}+2N_{M,t}}(p_{F,i}-p_{t-1}) + \frac{N_{F,b}}{N_{F,t}+2N_{M,t}}(p_{F,b}-p_{t-1}) \\
& + \frac{2N_{M,s}}{N_{F,t}+2N_{M,t}}(p_{M,s}-p_{t-1}) + \frac{2N_{M,i}}{N_{F,t}+2N_{M,t}}(p_{M,i}-p_{t-1}) + \frac{2N_{M,b}}{N_{F,t}+2N_{M,t}}(p_{M,b}-p_{t-1}) \Bigg] \\
= & \left(\frac{N_{F,s}}{N_{F,t}+2N_{M,t}}\right)^2 \mathrm{Var}(p_{F,s}-p_{t-1}) + \left(\frac{N_{F,i}}{N_{F,t}+2N_{M,t}}\right)^2 \mathrm{Var}(p_{F,i}-p_{t-1}) \\
& + \left(\frac{N_{F,b}}{N_{F,t}+2N_{M,t}}\right)^2 \mathrm{Var}(p_{F,b}-p_{t-1}) + \left(\frac{2N_{M,s}}{N_{F,t}+2N_{M,t}}\right)^2 \mathrm{Var}(p_{M,s}-p_{t-1}) \\
& + \left(\frac{2N_{M,i}}{N_{F,t}+2N_{M,t}}\right)^2 \mathrm{Var}(p_{M,i}-p_{t-1}) + \left(\frac{2N_{M,b}}{N_{F,t}+2N_{M,t}}\right)^2 \mathrm{Var}(p_{M,b}-p_{t-1}) \\
& + 2\frac{N_{F,s}2N_{M,s}}{(N_{F,t}+2N_{M,t})^2}\mathrm{Cov}(p_{F,s}-p_{t-1}, p_{M,s}-p_{t-1}) + 2\frac{N_{F,s}2N_{M,b}}{(N_{F,t}+2N_{M,t})^2}\mathrm{Cov}(p_{F,s}-p_{t-1}, p_{M,b}-p_{t-1}) \\
& + 2\frac{N_{F,i}2N_{M,i}}{(N_{F,t}+2N_{M,t})^2}\mathrm{Cov}(p_{F,i}-p_{t-1}, p_{M,i}-p_{t-1}) + 2\frac{N_{F,i}2N_{M,b}}{(N_{F,t}+2N_{M,t})^2}\mathrm{Cov}(p_{F,i}-p_{t-1}, p_{M,b}-p_{t-1}) \\
& + 2\frac{N_{F,b}2N_{M,s}}{(N_{F,t}+2N_{M,t})^2}\mathrm{Cov}(p_{F,b}-p_{t-1}, p_{M,s}-p_{t-1}) + 2\frac{N_{F,b}2N_{M,i}}{(N_{F,t}+2N_{M,t})^2}\mathrm{Cov}(p_{F,b}-p_{t-1}, p_{M,i}-p_{t-1}) \\
& + 2\frac{N_{F,b}2N_{M,b}}{(N_{F,t}+2N_{M,t})^2}\mathrm{Cov}(p_{F,b}-p_{t-1}, p_{M,b}-p_{t-1}) + 2\frac{2N_{M,s}2N_{M,b}}{(N_{F,t}+2N_{M,t})^2}\mathrm{Cov}(p_{M,s}-p_{t-1}, p_{M,b}-p_{t-1}) \\
& + 2\frac{2N_{M,i}2N_{M,b}}{(N_{F,t}+2N_{M,t})^2}\mathrm{Cov}(p_{M,i}-p_{t-1}, p_{M,b}-p_{t-1})
\end{aligned} \tag{14}$$

We assume immigrants in a given year are unrelated to survivors and set those covariances to zero. In addition, mothers cannot contribute a Z chromosome to daughters, so $\mathrm{Cov}\left(p_{F,s}-p_{t-1}, p_{F,b}-p_{t-1}\right)$ and $\mathrm{Cov}\left(p_{F,i}-p_{t-1}, p_{F,b}-p_{t-1}\right)$ are also zero.

#### Estimation of variance due to Mendelian segregation

For Z loci, only fathers can be heterozygous, as mothers have only one Z chromosome and are hemizygous. Here, we further partition the variance in allele frequency change due to births on the Z into variation due to varying family sizes and Mendelian segregation of alleles from heterozygous parents.

In year $t$, the allele frequency of the birth cohort at a given Z locus is the sum of the allele frequency ($g_{s,k}$) of each individual nestling $k$ of sex $s$. We will model this separately for males and females as follows:

$$p_{M,b} = \sum_{k=1}^{N_{M,b}} g_{M,k} \tag{15}$$

$$p_{F,b} = \sum_{k=1}^{N_{F,b}} g_{F,k} \tag{16}$$

$g_{M,k}$ has a value of 0, 0.5, or 1 for males with genotypes of 00, 01, or 11 respectively. $g_{F,k}$ has a value of 0 or 1 for females with genotypes of 0 or 1.

For males (*i.e.*, sons), we can write $g_{M,k}$ as:

$$g_{M,k} = \frac{1}{2}g_{k,dad} + \frac{1}{2}g_{k,mom} + \frac{1}{2}\delta_{k,dad} \tag{17}$$

where $g_{k,dad}$ and $g_{k,mom}$ are the allele frequencies of the father and mother of individual $k$. Each is multiplied by $\frac{1}{2}$ as each parent's contribution makes up one-half of the son's genotype. $\delta_{k,dad}$ is the difference in allele frequency between individual $k$ and his father due to Mendelian segregation; as a mother will always transmit her single Z to her son, there is no randomness and so no $\delta$ term for the mother.

The allele frequency of the male birth cohort in a given year is then:

$$\begin{aligned} p_{M,b} &= \sum_{k=1}^{N_{M,b}} \left[\frac{1}{2}(g_{k,dad} + g_{k,mom}) + \frac{1}{2}\delta_{k,dad}\right] \\ &= \frac{1}{2}(p_{M,dad} + p_{M,mom}) + \frac{1}{2}\sum_{k=1}^{N_{M,b}}(\delta_{k,dad}) \end{aligned} \tag{18}$$

where $p_{M,dad}$ is the allele frequency of the fathers of male nestlings born that year, weighted by the number of sons each father produced, and $p_{M,mom}$ is the allele frequency of the mothers of male nestlings born that year, similarly weighted by the number of sons each mother produced.

The allele frequency of the male birth cohort depends on both family size effects ($p_{M,b,fam}$) and Mendelian segregation of alleles in heterozygotes ($p_{M,b,mend}$). Thus, $p_{M,b}$ can be broken down into $p_{M,b,fam}$ and $p_{M,b,mend}$. We can now write $\text{Var}(p_{M,b} - p_{t-1})$ in terms of the variance due to family size and Mendelian noise:

$$\begin{aligned} \text{Var}(p_{M,b} - p_{t-1}) &= \text{Var}(p_{M,b,fam} + p_{M,b,mend} - p_{t-1}) \\ &= \text{Var}(p_{M,b,fam} - p_{t-1}) + \text{Var}(p_{M,b,mend}) \end{aligned} \tag{19}$$

To estimate the variance due to Mendelian noise, we solve for $p_{M,b,mend}$:

$$\text{Var}(p_{M,b,mend}) = \text{Var}\left[p_{M,b} - \frac{1}{2}\left(p_{M,dad} + p_{M,mom}\right)\right] \tag{20}$$

For females (*i.e.*, daughters), we can write $g_{F,k}$ as:

$$g_{F,k} = g_{k,dad} + \delta_{k,dad} \tag{21}$$

where $g_{k,dad}$ is the allele frequency of the father of individual $k$. Females inherit just one of their father's alleles as their single allele, and do not inherit a Z allele from their mother. Thus, unlike for sons, $g_{k,dad}$ is not multiplied by $\frac{1}{2}$ as the father's contribution makes up all of the daughter's genotype.

The allele frequency of the female birth cohort is then:

$$\begin{aligned} p_{F,b} &= \sum_{k=1}^{N_{F,b}} \left[g_{k,dad} + \delta_{k,dad}\right] \\ &= p_{F,dad} + \sum_{k=1}^{N_{F,b}}(\delta_{k,dad}) \end{aligned} \tag{22}$$

The first term is $p_{F,b,fam}$ and the second term is $p_{F,b,mend}$. We can now write $\text{Var}(p_{F,b} - p_{t-1})$ in terms of variance due to family size effects and Mendelian segregation:

$$\begin{aligned} \text{Var}(p_{F,b} - p_{t-1}) &= \text{Var}(p_{F,b,fam} + p_{F,b,mend} - p_{t-1}) \\ &= \text{Var}(p_{F,b,fam} - p_{t-1}) + \text{Var}(p_{F,b,mend}) \end{aligned} \tag{23}$$

The variance due to Mendelian noise is:

$$\text{Var}(p_{F,b,mend}) = \text{Var}\left[p_{F,b} - p_{F,dad}\right] \tag{24}$$

##### Sampling error

We correct for sampling error using the same approach we took for the autosomal model. Let $n_j$ be the number of genotyped individuals in a given category in a given year and $X_j$ be the observed allele counts. For males, let $x_j = \frac{X_j}{2n_j}$ be the observed allele frequency. For females, let $x_j = \frac{X_j}{n_j}$ be the observed allele frequency. The difference between the observed and true allele frequencies — the error due to sampling — may then be represented for both males and females by $\varepsilon_j$ such that $p_j = x_j - \varepsilon_j$.

The true allele frequencies $p_j$ in our model may now be broken down into observed allele frequencies and sampling error. Derivation for sampling error of Z-linked loci is identical to the autosomal model (see above for derivation), except for the Mendelian noise term. To incorporate error due to sampling in our estimate of Mendelian noise for males and females:

$$\begin{aligned}
\mathrm{Var}(p_{M,b,mend}) &= \mathrm{Var}\left[x_{M,b} - \varepsilon_{M,b} - \frac{1}{2}(x_{M,dad} - \varepsilon_{M,dad} + x_{M,mom} - \varepsilon_{M,mom})\right] \\
&= \mathrm{Var}\left[x_{M,b} - \frac{1}{2}\left(x_{M,dad} + x_{M,mom}\right)\right] - \mathrm{Var}\left[\varepsilon_{M,b} - \frac{1}{2}\left(\varepsilon_{M,dad} + \varepsilon_{M,mom}\right)\right] \qquad (25) \\
\mathrm{Var}(p_{M,b,fam} - p_{t-1}) &= \mathrm{Var}\left[\frac{1}{2}(p_{M,dad} + p_{M,mom}) - p_{t-1}\right] \\
&= \mathrm{Var}\left[\frac{1}{2}(x_{M,dad} - \varepsilon_{M,dad} + x_{M,mom} - \varepsilon_{M,mom}) - (x_{t-1} - \varepsilon_{t-1})\right] \\
&= \mathrm{Var}\left[\frac{1}{2}(x_{M,dad} + x_{M,mom}) - x_{t-1}\right] - \mathrm{Var}\left[\frac{1}{2}(\varepsilon_{M,dad} + \varepsilon_{M,mom}) - \varepsilon_{t-1}\right] \qquad (26)
\end{aligned}$$

$$\begin{aligned}
\mathrm{Var}(p_{F,b,mend}) &= \mathrm{Var}\left[(x_{F,b} - \varepsilon_{F,b}) - (x_{F,dad} - \varepsilon_{F,dad})\right] \\
&= \mathrm{Var}\left[x_{F,b} - x_{F,dad}\right] - \mathrm{Var}\left[\varepsilon_{F,b} - \varepsilon_{F,dad}\right] \qquad (27) \\
\mathrm{Var}(p_{F,b,fam} - p_{t-1}) &= \mathrm{Var}\left[p_{F,dad} - p_{t-1}\right] \\
&= \mathrm{Var}\left[(x_{F,dad} - \varepsilon_{F,dad}) - (x_{t-1} - \varepsilon_{t-1})\right] \\
&= \mathrm{Var}\left[x_{F,dad} - x_{t-1}\right] - \mathrm{Var}\left[\varepsilon_{F,dad} - \varepsilon_{t-1}\right] \qquad (28)
\end{aligned}$$

The covariance between the observed allele frequency and the error is again 0.

#### Allele frequency variance models with a fixed sex ratio

We used a similar approach to model allele frequency change while assuming a fixed sex ratio. Again basing our work off of the allele frequency model of [1], we divide each demographic group into separate components for males and females and relate the number of individuals in each category to the total number of individuals of that sex, rather than the total number of individuals in the whole population as above. Estimation of variance due to Mendelian segregation and sampling error is identical to previous sections.

##### Fixed sex ratio model for autosomal loci

For autosomal loci, let $N_{M,t}$ be the total number of males in the population in year $t$. Let $N_{M,s}$ be the number of males who survived from year $t-1$ to year $t$. Let $N_{M,i}$ be the number of males who immigrated into the population in year $t$. Let $N_{M,b}$ be the number of males born in the population in year $t$. All males in the population in a given year fit into one of these three categories (survivors, immigrants, or births) so $N_{M,t} = N_{M,s} + N_{M,i} + N_{M,b}$. If we represent the true allele counts for males in each category as $P_{M,j}$, then the true allele frequencies can be represented as $p_{M,j} = P_{M,j}/2N_{M,j}$ because each male has two copies of each autosome.

For females, let $N_{F,t}$ be the total number of females in the population in year $t$. Let $N_{F,s}$ be the number of females who survived from year $t-1$ to year $t$. Let $N_{F,i}$ be the number of females who immigrated into the population in year $t$. Let $N_{F,b}$ be the number of females born in the population in year $t$. All females in the population in a given year fit into one of these three categories (survivors, immigrants, or births) so $N_{F,t} = N_{F,s} + N_{F,i} + N_{F,b}$. If we represent the true allele counts for females in each category as $P_{F,j}$, then the true allele frequencies can be represented as $p_{F,j} = P_{F,j}/2N_{F,j}$ because each female has two copies of each autosome.

We assume a sex ratio of 0.5 on average, so 1/2 of the autosomes in the population are found in males and 1/2 of the autosomes in the population are found in females (*i.e.*, $P_t = \frac{1}{2}P_{M,t} + \frac{1}{2}P_{F,t}$). The change in allele frequencies for autosomal SNPs from year to year is:

$$
\begin{aligned}
\Delta p &= p_t - p_{t-1} \\
&= \frac{1}{2}p_{M,t} + \frac{1}{2}p_{F,t} - \frac{1}{2}p_{M,t-1} - \frac{1}{2}p_{F,t-1} \\
&= \frac{1}{2}\left[\frac{P_{M,t}}{2N_{M,t}} - \frac{P_{M,t-1}}{2N_{M,t-1}}\right] + \frac{1}{2}\left[\frac{P_{F,t}}{2N_{F,t}} - \frac{P_{F,t-1}}{2N_{F,t-1}}\right] \\
&= \frac{1}{2}\left[\frac{P_{M,s} + P_{M,i} + P_{M,b}}{2N_{M,t}}\frac{(N_{M,t-1})}{(N_{M,t-1})} - \frac{P_{M,t-1}}{2N_{M,t-1}}\frac{(N_{M,s} + N_{M,i} + N_{M,b})}{(N_{M,t})}\right] \\
&\quad + \frac{1}{2}\left[\frac{P_{F,s} + P_{F,i} + P_{F,b}}{2N_{F,t}}\frac{(N_{F,t-1})}{(N_{F,t-1})} - \frac{P_{F,t-1}}{2N_{F,t-1}}\frac{(N_{F,s} + N_{F,i} + N_{F,b})}{(N_{F,t})}\right] \\
&= \frac{1}{2}\left[\frac{P_{M,s}N_{M,t-1} - P_{M,t-1}N_{M,s} + P_{M,i}N_{M,t-1} - P_{M,t-1}N_{M,i} + P_{M,b}N_{M,t-1} - P_{M,t-1}N_{M,b}}{2N_{M,t}N_{M,t-1}}\right] \\
&\quad + \frac{1}{2}\left[\frac{P_{F,s}N_{F,t-1} - P_{F,t-1}N_{F,s} + P_{F,i}N_{F,t-1} - P_{F,t-1}N_{F,i} + P_{F,b}N_{F,t-1} - P_{F,t-1}N_{F,b}}{2N_{F,t}N_{F,t-1}}\right] \\
&= \frac{1}{2}\left[\frac{N_{M,s}}{N_{M,t}}\left(\frac{P_{M,s}}{2N_{M,s}} - \frac{P_{M,t-1}}{2N_{M,t-1}}\right) + \frac{N_{M,i}}{N_{M,t}}\left(\frac{P_{M,i}}{2N_{M,i}} - \frac{P_{M,t-1}}{2N_{M,t-1}}\right) + \frac{N_{M,b}}{N_{M,t}}\left(\frac{P_{M,b}}{2N_{M,b}} - \frac{P_{M,t-1}}{2N_{M,t-1}}\right)\right] \\
&\quad + \frac{1}{2}\left[\frac{N_{F,s}}{N_{F,t}}\left(\frac{P_{F,s}}{2N_{F,s}} - \frac{P_{F,t-1}}{2N_{F,t-1}}\right) + \frac{N_{F,i}}{N_{F,t}}\left(\frac{P_{F,i}}{2N_{F,i}} - \frac{P_{F,t-1}}{2N_{F,t-1}}\right) + \frac{N_{F,b}}{N_{F,t}}\left(\frac{P_{F,b}}{2N_{F,b}} - \frac{P_{F,t-1}}{2N_{F,t-1}}\right)\right] \\
&= \frac{1}{2}\left[\frac{N_{M,s}}{N_{M,t}}(p_{M,s} - p_{M,t-1}) + \frac{N_{M,i}}{N_{M,t}}(p_{M,i} - p_{M,t-1}) + \frac{N_{M,b}}{N_{M,t}}(p_{M,b} - p_{M,t-1})\right] \\
&\quad + \frac{1}{2}\left[\frac{N_{F,s}}{N_{F,t}}(p_{F,s} - p_{F,t-1}) + \frac{N_{F,i}}{N_{F,t}}(p_{F,i} - p_{F,t-1}) + \frac{N_{F,b}}{N_{F,t}}(p_{F,b} - p_{F,t-1})\right]
\end{aligned} \tag{29}
$$

We assume that immigrants in a given year are unrelated to survivors and to each other, so $\mathrm{Cov}(p_{M,s} - p_{M,t-1}, p_{M,i} - p_{M,t-1})$, $\mathrm{Cov}(p_{F,s} - p_{F,t-1}, p_{F,i} - p_{F,t-1})$, $\mathrm{Cov}(p_{M,s} - p_{M,t-1}, p_{F,i} - p_{F,t-1})$, $\mathrm{Cov}(p_{M,i} - p_{M,t-1}, p_{F,s} - p_{F,t-1})$, and $\mathrm{Cov}(p_{M,i} - p_{M,t-1}, p_{F,i} - p_{F,t-1})$ are all 0. The variance in allele frequency change over time is then:

$$
\begin{aligned}
\mathrm{Var}(\Delta p) = \mathrm{Var}\Bigg(&\frac{1}{2}\left[\frac{N_{M,s}}{N_{M,t}}(p_{M,s}-p_{M,t-1})+\frac{N_{M,i}}{N_{M,t}}(p_{M,i}-p_{M,t-1})+\frac{N_{M,b}}{N_{M,t}}(p_{M,b}-p_{M,t-1})\right]\\
&+\frac{1}{2}\left[\frac{N_{F,s}}{N_{F,t}}(p_{F,s}-p_{F,t-1})+\frac{N_{F,i}}{N_{F,t}}(p_{F,i}-p_{F,t-1})+\frac{N_{F,b}}{N_{F,t}}(p_{F,b}-p_{F,t-1})\right]\Bigg)\\
=&\left(\frac{N_{M,s}}{2N_{M,t}}\right)^2\mathrm{Var}(p_{M,s}-p_{M,t-1})+\left(\frac{N_{M,i}}{2N_{M,t}}\right)^2\mathrm{Var}(p_{M,i}-p_{M,t-1})\\
&+\left(\frac{N_{M,b}}{2N_{M,t}}\right)^2\mathrm{Var}(p_{M,b}-p_{M,t-1})+\left(\frac{N_{F,s}}{2N_{F,t}}\right)^2\mathrm{Var}(p_{F,s}-p_{F,t-1})\\
&+\left(\frac{N_{F,i}}{2N_{F,t}}\right)^2\mathrm{Var}(p_{F,i}-p_{F,t-1})+\left(\frac{N_{F,b}}{2N_{F,t}}\right)^2\mathrm{Var}(p_{F,b}-p_{F,t-1})\\
&+2\left(\frac{1}{2}\right)^2\left(\frac{N_{M,s}N_{M,b}}{N_{M,t}^2}\right)\mathrm{Cov}(p_{M,s}-p_{M,t-1},p_{M,b}-p_{M,t-1})\\
&+2\left(\frac{1}{2}\right)^2\left(\frac{N_{M,i}N_{M,b}}{N_{M,t}^2}\right)\mathrm{Cov}(p_{M,i}-p_{M,t-1},p_{M,b}-p_{M,t-1})\\
&+2\left(\frac{1}{2}\right)^2\left(\frac{N_{F,s}N_{F,b}}{N_{F,t}^2}\right)\mathrm{Cov}(p_{F,s}-p_{F,t-1},p_{F,b}-p_{F,t-1})\\
&+2\left(\frac{1}{2}\right)^2\left(\frac{N_{F,i}N_{F,b}}{N_{F,t}^2}\right)\mathrm{Cov}(p_{F,i}-p_{F,t-1},p_{F,b}-p_{F,t-1})\\
&+2\left(\frac{1}{2}\right)^2\left(\frac{N_{M,s}N_{F,s}}{N_{M,t}N_{F,t}}\right)\mathrm{Cov}(p_{M,s}-p_{M,t-1},p_{F,s}-p_{F,t-1})\\
&+2\left(\frac{1}{2}\right)^2\left(\frac{N_{M,s}N_{F,b}}{N_{M,t}N_{F,t}}\right)\mathrm{Cov}(p_{M,s}-p_{M,t-1},p_{F,b}-p_{F,t-1})\\
&+2\left(\frac{1}{2}\right)^2\left(\frac{N_{M,i}N_{F,b}}{N_{M,t}N_{F,t}}\right)\mathrm{Cov}(p_{M,i}-p_{M,t-1},p_{F,b}-p_{F,t-1})\\
&+2\left(\frac{1}{2}\right)^2\left(\frac{N_{M,b}N_{F,s}}{N_{M,t}N_{F,t}}\right)\mathrm{Cov}(p_{M,b}-p_{M,t-1},p_{F,s}-p_{F,t-1})\\
&+2\left(\frac{1}{2}\right)^2\left(\frac{N_{M,b}N_{F,i}}{N_{M,t}N_{F,t}}\right)\mathrm{Cov}(p_{M,b}-p_{M,t-1},p_{F,i}-p_{F,t-1})\\
&+2\left(\frac{1}{2}\right)^2\left(\frac{N_{M,b}N_{F,b}}{N_{M,t}N_{F,t}}\right)\mathrm{Cov}(p_{M,b}-p_{M,t-1},p_{F,b}-p_{F,t-1})
\end{aligned}
\tag{30}
$$

#### Fixed sex ratio model for Z loci

We use a similar approach to model allele frequency change for Z-linked loci, but again redefine a few terms and tweak our model slightly to account for the unique transmission rules of the Z chromosome. For males, let $N_{M,t}$ be the total number of males in the population in year $t$. Let $N_{M,s}$ be the number of males who survived from year $t-1$ to year $t$. Let $N_{M,i}$ be the number of males who immigrated into the population in year $t$. Let $N_{M,b}$ be the number of males born in the population in year $t$. All males in the population in a given year fit into one of these three categories (survivors, immigrants, or births) so $N_{M,t} = N_{M,s} + N_{M,i} + N_{M,b}$. If we represent the true allele counts for males in each category as $P_{M,j}$, then the true allele frequencies can be represented as $p_{M,j} = P_{M,j}/(2N_{M,j})$ because each male has two Z chromosomes.

Likewise, for females, let $N_{F,t}$ be the total number of females in the population in year $t$; $N_{F,s}$ the number of females who survived from year $t-1$ to year $t$; $N_{F,i}$ the number of females who immigrated into the population in year $t$; ans $N_{F,b}$ be the number of females born in the population in year $t$. As with males, $N_{F,t} = N_{F,s} + N_{F,i} + N_{F,b}$. If we represent the true allele counts for females in each category as $P_{F,j}$, then the true allele frequencies can be represented as $p_{F,j} = P_{F,j}/(N_{F,j})$ because each female has just one Z chromosome.

We assume a sex ratio of 0.5 on average, so 2/3 of the Z chromosomes in the population are found in males and 1/3 of the Z chromosomes in the population are found in females (*i.e.*, $P_t = \frac{2}{3}P_{M,t} + \frac{1}{3}P_{F,t}$). The empirical sex ratio is generally very close to 0.5 for every category except immigrants: 0.43-0.56 (mean 0.52) for survivors, 0.1-0.75 (mean 0.30) for immigrants, 0.42-0.59 (mean 0.51) for births, and 0.47-0.53 (mean 0.50) for the entire population.

The change in allele frequencies for SNPs on the Z chromosome (excluding the pseudoautosomal region) from year to year is:

$$\begin{aligned}
\Delta p &= p_t - p_{t-1} \\
&= \frac{2}{3}p_{M,t} + \frac{1}{3}p_{F,t} - \frac{2}{3}p_{M,t-1} - \frac{1}{3}p_{F,t-1} \\
&= \frac{2}{3}\left[\frac{P_{M,t}}{2N_{M,t}} - \frac{P_{M,t-1}}{2N_{M,t-1}}\right] + \frac{1}{3}\left[\frac{P_{F,t}}{N_{F,t}} - \frac{P_{F,t-1}}{N_{F,t-1}}\right] \\
&= \frac{2}{3}\left[\frac{P_{M,s}+P_{M,i}+P_{M,b}}{2N_{M,t}}\frac{(N_{M,t-1})}{(N_{M,t-1})} - \frac{P_{M,t-1}}{2N_{M,t-1}}\frac{(N_{M,s}+N_{M,i}+N_{M,b})}{(N_{M,t})}\right] \\
&\quad + \frac{1}{3}\left[\frac{P_{F,s}+P_{F,i}+P_{F,b}}{N_{F,t}}\frac{(N_{F,t-1})}{(N_{F,t-1})} - \frac{P_{F,t-1}}{N_{F,t-1}}\frac{(N_{F,s}+N_{F,i}+N_{F,b})}{(N_{F,t})}\right] \\
&= \frac{2}{3}\left[\frac{P_{M,s}N_{M,t-1} - P_{M,t-1}N_{M,s} + P_{M,i}N_{M,t-1} - P_{M,t-1}N_{M,i} + P_{M,b}N_{M,t-1} - P_{M,t-1}N_{M,b}}{2N_{M,t}N_{M,t-1}}\right] \\
&\quad + \frac{1}{3}\left[\frac{P_{F,s}N_{F,t-1} - P_{F,t-1}N_{F,s} + P_{F,i}N_{F,t-1} - P_{F,t-1}N_{F,i} + P_{F,b}N_{F,t-1} - P_{F,t-1}N_{F,b}}{N_{F,t}N_{F,t-1}}\right] \\
&= \frac{2}{3}\left[\frac{N_{M,s}}{N_{M,t}}\left(\frac{P_{M,s}}{2N_{M,s}} - \frac{P_{M,t-1}}{2N_{M,t-1}}\right) + \frac{N_{M,i}}{N_{M,t}}\left(\frac{P_{M,i}}{2N_{M,i}} - \frac{P_{M,t-1}}{2N_{M,t-1}}\right) + \frac{N_{M,b}}{N_{M,t}}\left(\frac{P_{M,b}}{2N_{M,b}} - \frac{P_{M,t-1}}{2N_{M,t-1}}\right)\right] \\
&\quad + \frac{1}{3}\left[\frac{N_{F,s}}{N_{F,t}}\left(\frac{P_{F,s}}{N_{F,s}} - \frac{P_{F,t-1}}{N_{F,t-1}}\right) + \frac{N_{F,i}}{N_{F,t}}\left(\frac{P_{F,i}}{N_{F,i}} - \frac{P_{F,t-1}}{N_{F,t-1}}\right) + \frac{N_{F,b}}{N_{F,t}}\left(\frac{P_{F,b}}{N_{F,b}} - \frac{P_{F,t-1}}{N_{F,t-1}}\right)\right] \\
&= \frac{2}{3}\left[\frac{N_{M,s}}{N_{M,t}}(p_{M,s} - p_{M,t-1}) + \frac{N_{M,i}}{N_{M,t}}(p_{M,i} - p_{M,t-1}) + \frac{N_{M,b}}{N_{M,t}}(p_{M,b} - p_{M,t-1})\right] \\
&\quad + \frac{1}{3}\left[\frac{N_{F,s}}{N_{F,t}}(p_{F,s} - p_{F,t-1}) + \frac{N_{F,i}}{N_{F,t}}(p_{F,i} - p_{F,t-1}) + \frac{N_{F,b}}{N_{F,t}}(p_{F,b} - p_{F,t-1})\right] \qquad (31)
\end{aligned}$$

The variance in allele frequency change over time is:

$$
\begin{aligned}
\mathrm{Var}(\Delta p) = \mathrm{Var}\Bigg(&\frac{2}{3}\left[\frac{N_{M,s}}{N_{M,t}}\left(p_{M,s}-p_{M,t-1}\right)+\frac{N_{M,i}}{N_{M,t}}\left(p_{M,i}-p_{M,t-1}\right)+\frac{N_{M,b}}{N_{M,t}}\left(p_{M,b}-p_{M,t-1}\right)\right] \\
&+\frac{1}{3}\left[\frac{N_{F,s}}{N_{F,t}}\left(p_{F,s}-p_{F,t-1}\right)+\frac{N_{F,i}}{N_{F,t}}\left(p_{F,i}-p_{F,t-1}\right)+\frac{N_{F,b}}{N_{F,t}}\left(p_{F,b}-p_{F,t-1}\right)\right]\Bigg) \\
= &\left(\frac{2N_{M,s}}{3N_{M,t}}\right)^2 \mathrm{Var}\left(p_{M,s}-p_{M,t-1}\right)+\left(\frac{2N_{M,i}}{3N_{M,t}}\right)^2 \mathrm{Var}\left(p_{M,i}-p_{M,t-1}\right) \\
&+\left(\frac{2N_{M,b}}{3N_{M,t}}\right)^2 \mathrm{Var}\left(p_{M,b}-p_{M,t-1}\right)+\left(\frac{N_{F,s}}{3N_{F,t}}\right)^2 \mathrm{Var}\left(p_{F,s}-p_{F,t-1}\right) \\
&+\left(\frac{N_{F,i}}{3N_{F,t}}\right)^2 \mathrm{Var}\left(p_{F,i}-p_{F,t-1}\right)+\left(\frac{N_{F,b}}{3N_{F,t}}\right)^2 \mathrm{Var}\left(p_{F,b}-p_{F,t-1}\right) \\
&+2\left(\frac{2}{3}\right)^2\left(\frac{N_{M,s}N_{M,b}}{N_{M,t}^2}\right)\mathrm{Cov}\left(p_{M,s}-p_{M,t-1},p_{M,b}-p_{M,t-1}\right) \\
&+2\left(\frac{2}{3}\right)^2\left(\frac{N_{M,i}N_{M,b}}{N_{M,t}^2}\right)\mathrm{Cov}\left(p_{M,i}-p_{M,t-1},p_{M,b}-p_{M,t-1}\right) \\
&+2\left(\frac{2}{3}\right)\left(\frac{1}{3}\right)\left(\frac{N_{M,s}N_{F,s}}{N_{M,t}N_{F,t}}\right)\mathrm{Cov}\left(p_{M,s}-p_{M,t-1},p_{F,s}-p_{F,t-1}\right) \\
&+2\left(\frac{2}{3}\right)\left(\frac{1}{3}\right)\left(\frac{N_{M,s}N_{F,b}}{N_{M,t}N_{F,t}}\right)\mathrm{Cov}\left(p_{M,s}-p_{M,t-1},p_{F,b}-p_{F,t-1}\right) \\
&+2\left(\frac{2}{3}\right)\left(\frac{1}{3}\right)\left(\frac{N_{M,i}N_{F,b}}{N_{M,t}N_{F,t}}\right)\mathrm{Cov}\left(p_{M,i}-p_{M,t-1},p_{F,b}-p_{F,t-1}\right) \\
&+2\left(\frac{2}{3}\right)\left(\frac{1}{3}\right)\left(\frac{N_{M,b}N_{F,s}}{N_{M,t}N_{F,t}}\right)\mathrm{Cov}\left(p_{M,b}-p_{M,t-1},p_{F,s}-p_{F,t-1}\right) \\
&+2\left(\frac{2}{3}\right)\left(\frac{1}{3}\right)\left(\frac{N_{M,b}N_{F,i}}{N_{M,t}N_{F,t}}\right)\mathrm{Cov}\left(p_{M,b}-p_{M,t-1},p_{F,i}-p_{F,t-1}\right) \\
&+2\left(\frac{2}{3}\right)\left(\frac{1}{3}\right)\left(\frac{N_{M,b}N_{F,b}}{N_{M,t}N_{F,t}}\right)\mathrm{Cov}\left(p_{M,b}-p_{M,t-1},p_{F,b}-p_{F,t-1}\right)
\end{aligned} \tag{32}
$$

Recall that covariances between immigrants and survivors are zero because we assume immigrants in a given year are unrelated to survivors, and $\mathrm{Cov}\left(p_{F,s}-p_{t-1},p_{F,b}-p_{t-1}\right)$ and $\mathrm{Cov}\left(p_{F,i}-p_{t-1},p_{F,b}-p_{t-1}\right)$ are zero because mothers cannot contribute a Z chromosome to daughters.

### Supplementary Material

#### Genome assembly and annotation

Here, we generated an improved version (v2) of the Florida Scrub-Jay genome by performing additional scaffolding. The first version of the Florida Scrub-Jay genome (GenBank assembly GCA_013398375.1; [2]) was assembled from Illumina data generated from a male (ZZ) using ALLPATHS-LG ([3]). Dovetail Genomics created Chicago and Dovetail Hi-C libraries from blood samples taken from the same individual sequenced for the Illumina-only assembly, generating 330 million and 324 million 2x150 bp reads (99 GB and 97.5 GB) respectively, then scaffolded the genome using their HiRise pipeline. We then used Chromonomer v1.13 ([4]) and Juicer v.1.5.6 ([5]) along with a preliminary framework linkage map built using CRIMAP 2.507 ([6]; see Romero *et al. in prep.* for details) to confirm the assembly. This new assembly is 1.06 Gb in size, with 878 scaffolds and a scaffold N50 of 74.35 Mb. While no karyotype data exist for the Florida Scrub-Jay, a typical avian genome has $2n = 76$–80 chromosomes, with a few macrochromosomes and several microchromosomes ([7]). [8] use a karyotype of $2n = 80$ for the closely-related California Scrub-Jay (*Aphelocoma californica*) based on the median chromosome number for other members of the Corvidae family. Based on sequence homology with the zebra finch (*Taeniopygia guttata*) genome assembly version 3.2.4, we putatively assigned chromosome identities to 31 scaffolds (Table S1). We ran BUSCO v.5.2.2 ([9]) using the Aves ortholog database (aves_odb10) to evaluate the quality of our genome assembly and obtained a BUSCO completeness score of 98%.

For annotation, we first performed *de novo* repeat annotation using Repeatmodeler v.2.0.1. We then concatenated our repeat library with the repeat library from ([10]) to soft-mask our genome using Repeatmasker v.4.1.0. Next, we ran BRAKER2 v.2.1.6 ([11]) on the soft-masked genome using default parameters, seeding the protein annotation with the vertebrate OrthoDB database and all chicken genes from UniProt. We then ran BRAKER1 v.2.1.6 ([12]) and StringTie ([13]) using RNA-seq data generated from tissues collected by ([14]). We used Qiagen RNeasy kits to extract RNA from liver, heart, and kidney samples from one male and one female, in addition to ovary from the female, and performed two lanes of 2x101 bp Illumina HiSeq sequencing. We then combined the output from all three annotations using a custom R script. We ran BUSCO v.5.2.2 ([9]) on our final annotation. Our annotation resulted in 21,958 genes, representing a BUSCO score of 92%.

To determine the physical locations of the Beadchip SNPs based on the version 2 assembly, we aligned the Beadchip probe sequences to the new reference assembly using bwa ([15]) and used the alignment results to calculate updated SNP locations. Our SNPs are located on 46 different scaffolds and 31 putative chromosomes (Table S1). We used SnpEff v4.3 ([16]) to perform variant annotation. For variants with multiple annotations, we report the annotation with the highest putative effect. We found that most of our SNPs are intergenic (68.6%) or intragenic/intronic (25.9%). Only 5.4% of sites are predicted to be in exons; 2.3% are synonymous and 3.1% are nonsynonymous.

#### The pseudoautosomal region

We putatively identified the pseudoautosomal region (PAR) based on a preliminary framework linkage map built using CRIMAP 2.507 ([6]; see Romero *et al. in prep.* for details). We identified a few SNPs from scaffolds 3451 and 439 of the version 1 genome assembly that were segregating autosomally, and assigned all SNPs on those scaffolds or more distal to the PAR. Given the sparseness of our preliminary framework map, we cannot define the exact boundary of the PAR. Our rough estimate is that the PAR is at least 0.66 Mb in length.

We do not include the PAR in our analysis of expected genetic contributions on the Z; expected genetic contribution simulations for the Z were performed using the transmission rules for Z-linked loci not in the PAR. We also do not include the PAR in our allele frequency variance models. Given the unique evolutionary trajectory of PAR alleles ([17]), we included the 19 PAR SNPs in our tests of selection, using autosomal transmission rules. We found no sites with significant evidence of selection among the 19 PAR SNPs.

#### Unsexed individuals and expected genetic contributions

Our population pedigree contains 1,896 unsexed individuals. None of these birds produced any offspring, so unsexed individuals only impact expected genetic contributions to their natal year. As most unsexed individuals were born before 2005, and we are primarily interested in expected genetic contributions to more recent years, our approach for assigning sexes to these individuals does not significantly impact our results. To demonstrate, we estimated expected genetic contributions of 926 breeders using different sex ratios for assigning sexes. Expected genetic contributions to the population in 2013 for both autosomes and the Z chromosome are nearly identical if we assign all unsexed individuals as females or males (Spearman's $\rho >$ 0.99 and $p <$2.2e-16 for both; Fig. S1) - the slight differences in expected Z genetic contributions are caused by the 4 unsexed 2013 nestlings.

#### Expected genetic contributions of male and female partners

We explored the relationship between genealogical and expected genetic contributions for autosomes and the Z chromosome by comparing the contributions of male and female partners. While Florida Scrub-Jays are monogamous, many individuals pair again if their partner dies. For the small subset of pairs that only bred with each other throughout their entire life (*i.e.*, lifelong monogamous pairs), the male and female partners always had equal expected autosomal genetic contributions in 2013 (Fig. S4A), but some males had higher expected Z genetic contributions than their mate, especially when the sex ratio of their offspring was skewed towards female offspring (Fig. S4B). Recall that mothers only transmit their Z chromosomes to their sons and therefore do not contribute Z alleles to their daughters or any descendants of their daughters. We examined the pedigrees of all descendants for the six lifelong monogamous pairs with unequal expected Z genetic contributions between partners. The three female breeders with zero expected Z genetic contributions in 2013 had only daughters that contributed to the 2013 birth cohort, while the three female breeders with greater than zero expected Z genetic contributions had descendants in the 2013 birth cohort via both sons and daughters. The fact that Florida Scrub-Jays can have multiple mates throughout their lifetime may increase the difference in expected genetic contributions between male-female pairs. To confirm this hypothesis, we compared the expected genetic contributions of all male and female partners and tracked the number of mates each individual had throughout their lifetime (Fig. S4C and D). For breeding pairs in which the female breeder had more mates throughout her lifetime compared to the male breeder, the female tended to have higher expected autosomal genetic contributions than her mate, and the opposite was true if the male breeder had a higher number of mates. We found a similar pattern for the Z chromosome. However, many females had lower expected Z genetic contributions than their male partners despite having a higher number of mates because of the important role the sex ratio of descendants plays in determining expected Z genetic contributions.

#### Allele frequency variance model implementation

We modeled the variance in allele frequency change over time using all individuals present in a core set of territories in 1990-2013. Individuals present in 1990 were assigned as founders; in subsequent years, individuals were assigned as male or female surviors, immigrants, or nestlings (births). Figure S2 shows the number of individuals of each sex and the proportion genotyped in each demographic group for each year.

We calculated allele frequencies among genotyped individuals for each category in each year at 10,731 autosomal SNPs for the autosomal model and 250 Z-linked SNPs for the Z chromosome model. We then empirically estimated sampling error by assigning genotypes to the founders present in 1990 based on the observed allele frequency spectrum for the autosomal or Z-linked loci, respectively, then simulating Mendelian transmission forward in time for 100,000 replicates. For the Z, this simulation requires that every individual be sexed, so we randomly assigned unsexed individuals as males or females based on the empirical sex ratio (which is approximately 50:50). We averaged across all loci to obtain the variance in allele frequency change contributed by each term. Due to low genotyping rates in 1990-1998, we consider only the period of 1999-2013 in our results.

Allele frequency variance models were implemented in R v. 4.0.2 and R v. 3.6.3 ([18]) with packages base, stats, foreach ([19]), doParallel ([20]), plyr ([21]), and dplyr ([22]) and visualized with packages ggplot2 ([23]) and cowplot ([24]).

#### Allele frequency variance models with a fixed sex ratio

We observed a negative covariance between male and female survivors, which could be a mathematical artifact arising because survivors form a large proportion of the population from the previous year and therefore have a similar mean allele frequency to the total allele frequency from last year ($p_{t-1}$). When the group of survivors is subdivided into males and females, the mean allele frequency of one group will, by chance, be above $p_{t-1}$, and the mean of the other group below $p_{t-1}$, creating a negative covariance between male and female survivors. To test this hypothesis, we ran an additional set of models that assumed a 50:50 sex ratio for each demographic group (survivors, immigrants, and births). This assumption is fairly reasonable for survivors and births, but not for immigrants (Fig. S2). By assuming a 50:50 sex ratio in our additional set of models, we were able to compare the allele frequency of each category to the allele frequency of individuals of that sex the previous year (*i.e.*, $p_{M,t-1}$ or $p_{F,t-1}$) and avoid the artifact described above. The equations for the change in allele frequencies between two years for an autosomal locus and for a Z-linked locus assuming a 50:50 sex ratio are derived in Appendix A. These models did not have the same biases based on properties of the mean, and indeed we found near-zero covariances between male and female survivors (Fig. S11).

#### Allele frequency variance models with LD-pruning

Variation in recombination rates across the genome may influence our estimates of the variance in allele frequency change over time. To assess the impact of linkage on our model results, we re-ran the allele frequency variance analysis using a set of LD-pruned SNPs. We used the 'indep-pairwise' function in PLINK [25] to remove SNPS with an $r^2 > 0.1$ using a window size of 50 kb and variant step size of 5. Our LD-pruned dataset consisted of 4,200 SNPs. The variance component estimates we obtained with our full dataset and the LD-pruned dataset are highly correlated (Spearman's $\rho = 0.99$ for autosomes and 0.94 for the Z chromosome, $p$ <2.2e-16 for both; Fig. S9). Confidence intervals for the estimates obtained from the pruned dataset are larger than those from the full dataset for both autosomes (mean CI = 0.25 for the full dataset and 0.30 for the pruned dataset; t = 24.538, df = 237, $p$ <2.2e-16) and the Z chromosome (mean CI = 0.33 for the full dataset and 0.41 for the pruned dataset; t = 18.133, df = 209, $p$ <2.2e-16).

### Supplementary Tables

| Chromosome | Size | # SNPs |
|---|---|---|
| 1 | 118618941 | 700 |
| 1A | 74350943 | 486 |
| 2 | 156027883 | 766 |
| 3 | 116014166 | 709 |
| 4 | 73917176 | 369 |
| 4A | 20491957 | 329 |
| 5 | 64039664 | 539 |
| 6 | 36332134 | 349 |
| 7 | 39276876 | 369 |
| 8 | 31891339 | 393 |
| 9 | 26024946 | 364 |
| 10 | 20796167 | 304 |
| 11 | 21126892 | 264 |
| 12 | 21104894 | 313 |
| 13 | 19131323 | 414 |
| 14 | 16732657 | 467 |
| 15 | 14520692 | 310 |
| 17 | 11119423 | 374 |
| 18 | 12207174 | 341 |
| 19 | 11192798 | 310 |
| 20 | 15072428 | 363 |
| 21 | 8410644 | 201 |
| 22 | 5244962 | 165 |
| 23 | 7401491 | 278 |
| 24 | 6926109 | 265 |
| 25 | 3360146 | 98 |
| 26 | 6773329 | 319 |
| 27 | 5848943 | 222 |
| 28 | 5850724 | 211 |
| LGE22 | 8882011 | 113 |
| Z | 75605511 | 269 |
| Unmapped | 6675375 | 26 |

**Table S1:** Chromosome sizes from the version 2 Florida Scrub-Jay genome assembly and the number of Beadchip SNPs on each chromosome. We assigned chromosome names to individual scaffolds based on sequence homology to the Zebra Finch genome. There are 847 unmapped scaffolds, 15 of which have at least one Beadchip SNP.

| Chromosome | Term | Estimate | S.E. | $t$ | $p$ |
|---|---|---|---|---|---|
| Autosome | Intercept | $3.49\times10^{-4}$ | $9.38\times10^{-5}$ | 3.72 | $2.11\times10^{-4}$*** |
| | Genealogical Contribution | 0.070 | 0.002 | 30.37 | $<2.2\times10^{-16}$*** |
| | Sex | $2.94\times10^{-5}$ | $1.30\times10^{-4}$ | 0.23 | 0.82 |
| | Genealogical Contribution × Sex | -0.004 | 0.003 | -1.39 | 0.16 |
| Z | Intercept | $6.32\times10^{-4}$ | $1.31\times10^{-4}$ | 4.83 | $1.62\times10^{-6}$*** |
| | Genealogical Contribution | 0.083 | 0.003 | 25.81 | $<2.2\times10^{-16}$*** |
| | Sex | $-2.04\times10^{-4}$ | $1.82\times10^{-4}$ | -1.12 | 0.26 |
| | Genealogical Contribution × Sex | -0.043 | 0.004 | -9.62 | $<2.2\times10^{-16}$*** |

**Table S2:** Linear model results for autosomal or Z-linked expected individual genetic contributions to the population in 2013 as a function of genealogical contributions and sex. Degrees of freedom = 922 and multiple $R^2 = 0.66$ for both models. Significant estimates are indicated as follows: *$p < 0.05$, **$p < 0.01$, ***$p < 0.001$.

| Coefficient | Estimate | S.E. | $t$ | $p$ |
|---|---|---|---|---|
| Autosomal expected contributions | 1.24 | 0.024 | 51.52 | $<2.2\times10^{-16}$*** |
| Female | $1.33\times10^{-4}$ | $8.41\times10^{-5}$ | 1.58 | 0.12 |
| Male | $1.12\times10^{-4}$ | $8.13\times10^{-5}$ | 1.38 | 0.17 |
| Autosomal expected contributions:Male | -0.57 | 0.034 | -16.89 | $<2.2\times10^{-16}$*** |

**Table S3:** Linear model results for Z expected genetic contributions to the population in 2013 as a function of autosomal expected genetic contributions and sex. Degrees of freedom = 922 and multiple $R^2 = 0.82$. Significant estimates are indicated as follows: *$p < 0.05$, **$p < 0.01$, ***$p < 0.001$.

| Coefficient | Estimate | S.E. | $t$ | $p$ |
|---|---|---|---|---|
| Intercept | 8.27 | 0.91 | 9.10 | $1.03\times10^{-15}$*** |
| Sex | 2.33 | 0.49 | 4.79 | $4.30\times10^{-6}$*** |
| ImmCohort | -0.41 | 0.06 | -6.83 | $2.61\times10^{-10}$*** |

**Table S4:** Linear model results for the number of incoming immigrants as a function of sex and cohort. Degrees of freedom = 135 and multiple $R^2 = 0.34$. Significant estimates are indicated as follows: *$p < 0.05$, **$p < 0.01$, ***$p < 0.001$.

| Chromosome | Coefficient | Estimate | S.E. | $t$ | $p$ |
|---|---|---|---|---|---|
| Autosome | Intercept | -0.012 | 0.014 | -0.89 | 0.39 |
| | Immigrant cohort size | 0.002 | 0.0008 | 3.00 | 0.012* |
| | Sex (male) | 0.007 | 0.007 | 0.99 | 0.34 |
| Z | Intercept | -0.008 | 0.016 | -0.50 | 0.63 |
| | Immigrant cohort size | 0.002 | 0.001 | 1.69 | 0.12 |
| | Sex (male) | 0.019 | 0.008 | 2.32 | 0.04* |

**Table S5:** Linear model results for levels of autosomal or Z-linked gene flow as a function of sex and immigrant cohort size. Degrees of freedom = 11 for both models, and multiple $R^2 = 0.45$ for autosomes and 0.36 for the Z. Significant estimates are indicated as follows: *$p < 0.05$, **$p < 0.01$, ***$p < 0.001$.

| SNP | Position | Annotation | Year | Obs. $\Delta p$ | Sim. $\Delta p$ | $p$ | $q$ |
|---|---|---|---|---|---|---|---|
| 12182 | 331669 | intergenic region | 1999-2013 | 0.043 | -0.166 | $1.85\times10^{-4}$ | 0.213 |
| 11931 | 33695922 | intergenic region | 1999-2000 | 0.024 | -0.017 | $1\times10^{-6}$ | 0.000* |
| 12023 | 72133876 | intergenic region | 1999-2000 | -0.039 | -0.039 | $1\times10^{-6}$ | 0.000* |
| 12177 | 482246 | upstream gene variant | 2000-2001 | -0.008 | -0.005 | $1\times10^{-6}$ | 0.000* |
| 12085 | 66242929 | downstream gene variant | 2000-2001 | 0.003 | 0.000 | $1\times10^{-6}$ | 0.000* |
| 12023 | 72133876 | intergenic region | 2000-2001 | 0.027 | 0.027 | $1\times10^{-6}$ | 0.000* |
| 11959 | 20975141 | intron variant | 2001-2002 | -0.009 | -0.047 | $1\times10^{-6}$ | 0.000* |

**Table S6:** Z-linked SNPs with significant ($q < 0.25$) net allele frequency shifts in 1999-2013 or between adjacent years in 1999-2013. SNP position is from the Florida Scrub-Jay genome v2, and variant annotation is from SnpEff. $\Delta p$ = change in allele frequency (observed or simulated); $p$ = p-value; $q$ = adjusted p-value (False Discovery Rate correction) for multiple comparisons across all SNPs in the genome (n = 11,000). SNPs that are significant at a more stringent FDR cutoff of $q < 0.1$ are marked with an *.

| SNP | Chr | Position | Annotation | Obs. $\Delta p$ | Sim. $\Delta p$ | 2019 $q$ | $p$ | $q$ |
|---|---|---|---|---|---|---|---|---|
| 11034 | 1A | 69547460 | intergenic region | -0.256 | -0.135 | 0.220 | $3.06\times10^{-4}$ | 0.213 |
| 2181 | 3 | 13362610 | intergenic region | 0.150 | 0.029 | 0.220 | $4.20\times10^{-5}$ | 0.213 |
| 2231 | 3 | 16445530 | intergenic region | -0.116 | 0.008 | 0.220 | $1.69\times10^{-4}$ | 0.213 |
| 3123 | 4 | 5126617 | downstream gene variant | -0.031 | 0.087 | 0.243 | $4.09\times10^{-4}$ | 0.243 |
| 3056 | 4 | 16020259 | intergenic region | 0.089 | 0.008 | 0.220 | $2.72\times10^{-4}$ | 0.213 |
| 3602 | 5 | 5869549 | upstream gene variant | -0.124 | -0.036 | 0.220 | $2.59\times10^{-4}$ | 0.213 |
| 3955 | 6 | 13793244 | intron variant | -0.172 | -0.041 | 0.220 | $2.79\times10^{-4}$ | 0.213 |
| 3950 | 6 | 13842442 | intron variant | -0.139 | -0.015 | 0.220 | $3.15\times10^{-4}$ | 0.213 |
| 3791 | 6 | 32608847 | intron variant | 0.078 | 0.011 | 0.220 | $2.23\times10^{-4}$ | 0.213 |
| 4924 | 8 | 2993342 | intergenic region | 0.149 | -0.003 | 0.220 | $2.75\times10^{-5}$ | 0.213 |
| 4578 | 8 | 31424463 | downstream gene variant | -0.090 | -0.010 | 0.220 | $2.84\times10^{-4}$ | 0.213 |
| 6389 | 12 | 970548 | downstream gene variant | -0.038 | 0.076 | 0.240 | $3.82\times10^{-4}$ | 0.240 |
| 9045 | 20 | 9981163 | intergenic region | -0.089 | 0.022 | 0.220 | $2.97\times10^{-4}$ | 0.213 |
| 9028 | 20 | 10283362 | intergenic region | -0.086 | 0.012 | 0.220 | $2.42\times10^{-4}$ | 0.213 |
| 8908 | 20 | 14090934 | upstream gene variant | 0.052 | -0.071 | 0.220 | $2.81\times10^{-4}$ | 0.213 |
| 10255 | 24 | 893755 | intron variant | -0.022 | 0.096 | 0.220 | $1.77\times10^{-4}$ | 0.213 |
| 10320 | 25 | 3198383 | upstream gene variant | 0.055 | -0.055 | 0.220 | $3.29\times10^{-4}$ | 0.213 |
| 273 | 26 | 3308218 | upstream gene variant | 0.136 | 0.022 | 0.220 | $2.72\times10^{-4}$ | 0.213 |

**Table S7:** Autosomal SNPs with a significant ($q < 0.25$) net allele frequency shift in 1999-2013 as detected by [1], reassessed to correct for the larger number of comparisons after inclusion of the Z-linked SNPs and allow for pseudocounts. SNP position is from the Florida Scrub-Jay genome v2, and variant annotation is from SnpEff. $\Delta p$ = change in allele frequency (observed or simulated); 2019 $q$ = adjusted p-value (False Discovery Rate correction) from [1] for multiple comparisons across autosomal SNPs (n = 10,731); $p$ = updated p-value with pseudocounts included; $q$ = adjusted p-value for multiple comparisons across all SNPs in the genome (n = 11,000). No SNPs are significant at a more stringent FDR cutoff of $q < 0.1$.

| SNP | Chr | Position | Annotation | Year | Obs. $\Delta p$ | Sim. $\Delta p$ | 2019 $q$ | $p$ | $q$ |
|---|---|---|---|---|---|---|---|---|---|
| 6873 | 14 | 663946 | intergenic region | 1999-2000 | 0.178 | 0.087 | 0.350 | $6.10 \times 10^{-5}$ | 0.224 |
| 1055 | 1 | 1498756 | upstream gene variant | 2000-2001 | 0.027 | -0.060 | 0.319 | $9.50 \times 10^{-5}$ | 0.209 |
| 1771 | 2 | 30481558 | intergenic region | 2000-2001 | 0.063 | -0.008 | 0.319 | $7.65 \times 10^{-5}$ | 0.209 |
| 9167 | 20 | 6610582 | intron variant | 2000-2001 | -0.049 | 0.000 | 0.319 | $1.36 \times 10^{-4}$ | 0.248 |
| 10963 | 1A | 72543265 | intron variant | 2001-2002 | -0.061 | -0.004 | 0.168 | $2.05 \times 10^{-4}$ | 0.161 |
| 1774 | 2 | 30123409 | intergenic region | 2001-2002 | 0.096 | 0.016 | 0.160 | $1.41 \times 10^{-4}$ | 0.151 |
| 1771 | 2 | 30481558 | intergenic region | 2001-2002 | -0.065 | 0.014 | 0.052* | $1.55 \times 10^{-5}$ | 0.043* |
| 2679 | 3 | 109275367 | intergenic region | 2001-2002 | -0.063 | 0.011 | 0.035* | $5.50 \times 10^{-6}$ | 0.028* |
| 2757 | 4 | 287107 | intergenic region | 2001-2002 | 0.034 | -0.003 | 0.089* | $5.05 \times 10^{-5}$ | 0.079* |
| 11549 | 4A | 5244498 | upstream gene variant | 2001-2002 | 0.072 | -0.001 | 0.249 | $3.27 \times 10^{-4}$ | 0.239 |
| 4139 | 6 | 1027351 | intergenic region | 2001-2002 | -0.123 | -0.062 | 0.160 | $1.30 \times 10^{-4}$ | 0.151 |
| 4212 | 7 | 38876364 | upstream gene variant | 2001-2002 | -0.167 | -0.071 | 0.035* | $7.50 \times 10^{-6}$ | 0.028* |
| 4199 | 7 | 39041415 | missense variant | 2001-2002 | 0.051 | -0.044 | 0.089* | $4.70 \times 10^{-5}$ | 0.079* |
| 6730 | 13 | 6717300 | upstream gene variant | 2001-2002 | -0.052 | 0.033 | 0.160 | $1.52 \times 10^{-4}$ | 0.151 |
| 9167 | 20 | 6610582 | intron variant | 2001-2002 | 0.067 | 0.016 | 0.089* | $3.80 \times 10^{-5}$ | 0.079* |
| 8947 | 20 | 13186622 | intergenic region | 2001-2002 | -0.060 | 0.036 | 0.168 | $2.01 \times 10^{-4}$ | 0.161 |
| 9463 | 21 | 432433 | intron variant | 2001-2002 | 0.138 | 0.045 | 0.101 | $6.70 \times 10^{-5}$ | 0.092* |
| 10760 | 28 | 3732531 | upstream gene variant | 2001-2002 | 0.026 | -0.040 | 0.160 | $1.65 \times 10^{-4}$ | 0.151 |
| 3003 | 4 | 39075056 | intergenic region | 2003-2004 | -0.108 | -0.002 | 0.247 | $7.00 \times 10^{-5}$ | 0.257 |
| 4648 | 8 | 29961336 | upstream gene variant | 2003-2004 | -0.147 | -0.041 | 0.247 | $3.50 \times 10^{-5}$ | 0.257 |
| 8146 | 17 | 1015618 | upstream gene variant | 2003-2004 | 0.074 | -0.031 | 0.247 | $4.75 \times 10^{-5}$ | 0.257 |
| 1660 | 2 | 61330851 | splice region variant | 2005-2006 | 0.115 | 0.006 | 0.185 | $7.15 \times 10^{-5}$ | 0.191 |
| 4442 | 7 | 30284642 | intergenic region | 2005-2006 | 0.071 | -0.010 | 0.185 | $7.20 \times 10^{-5}$ | 0.191 |
| 4447 | 7 | 30768473 | intergenic region | 2005-2006 | 0.065 | -0.004 | 0.185 | $1.22 \times 10^{-4}$ | 0.191 |
| 8256 | 18 | 1799320 | intron variant | 2005-2006 | -0.092 | 0.012 | 0.185 | $1.11 \times 10^{-4}$ | 0.191 |
| 8272 | 18 | 2451604 | intron variant | 2005-2006 | -0.078 | -0.007 | 0.185 | $9.40 \times 10^{-5}$ | 0.191 |
| 8532 | 18 | 5127999 | intergenic region | 2005-2006 | -0.109 | 0.008 | 0.185 | $5.40 \times 10^{-5}$ | 0.191 |
| 8344 | 18 | 11035451 | intergenic region | 2005-2006 | 0.129 | 0.016 | 0.185 | $9.15 \times 10^{-5}$ | 0.191 |
| 2548 | 3 | 78929023 | intergenic region | 2006-2007 | -0.180 | -0.052 | 0.182 | $1.80 \times 10^{-5}$ | 0.198 |
| 1754 | 2 | 35522200 | intron variant | 2009-2010 | -0.104 | -0.012 | 0.232 | $2.24 \times 10^{-4}$ | 0.239 |
| 2440 | 3 | 50364194 | intergenic region | 2009-2010 | -0.067 | -0.006 | 0.239 | $3.02 \times 10^{-4}$ | 0.246 |
| 2989 | 4 | 45250978 | upstream gene variant | 2009-2010 | -0.088 | 0.059 | 0.232 | $1.76 \times 10^{-4}$ | 0.239 |
| 3655 | 5 | 3948756 | upstream gene variant | 2009-2010 | -0.132 | 0.016 | 0.232 | $1.97 \times 10^{-4}$ | 0.239 |
| 3510 | 5 | 12201294 | upstream gene variant | 2009-2010 | -0.078 | 0.050 | 0.232 | $1.18 \times 10^{-4}$ | 0.239 |
| 3178 | 5 | 61210587 | downstream gene variant | 2009-2010 | -0.073 | -0.021 | 0.232 | $1.37 \times 10^{-4}$ | 0.239 |
| 3195 | 5 | 62364412 | upstream gene variant | 2009-2010 | 0.081 | -0.047 | 0.239 | $3.13 \times 10^{-4}$ | 0.246 |
| 4956 | 8 | 1544848 | downstream gene variant | 2009-2010 | -0.071 | -0.016 | 0.232 | $1.55 \times 10^{-4}$ | 0.239 |

| | | | | | | | | | |
|---|---|---|---|---|---|---|---|---|---|
| 4704 | 8 | 24098695 | intergenic region | 2009-2010 | -0.095 | 0.039 | 0.233 | $2.62 \times 10^{-4}$ | 0.240 |
| 4691 | 8 | 24412799 | intergenic region | 2009-2010 | -0.082 | 0.042 | 0.232 | $1.13 \times 10^{-4}$ | 0.239 |
| 6737 | 13 | 6761655 | upstream gene variant | 2009-2010 | -0.158 | -0.003 | 0.232 | $2.40 \times 10^{-5}$ | 0.239 |
| 10258 | 24 | 872600 | upstream gene variant | 2009-2010 | 0.058 | -0.031 | 0.232 | $1.75 \times 10^{-4}$ | 0.239 |
| 10242 | 24 | 1049980 | downstream gene variant | 2009-2010 | 0.109 | 0.002 | 0.232 | $1.42 \times 10^{-4}$ | 0.239 |
| 181 | 26 | 1620253 | upstream gene variant | 2009-2010 | -0.117 | -0.002 | 0.232 | $2.39 \times 10^{-4}$ | 0.239 |
| 1782 | 2 | 29280160 | intergenic region | 2010-2011 | -0.122 | -0.002 | 0.054* | $6.00 \times 10^{-6}$ | 0.066* |
| 1644 | 2 | 64208514 | intergenic region | 2011-2012 | 0.167 | 0.010 | 0.236 | $9.55 \times 10^{-5}$ | 0.244 |
| 1322 | 2 | 146541682 | intergenic region | 2011-2012 | 0.101 | 0.018 | 0.236 | $7.10 \times 10^{-5}$ | 0.244 |
| 11790 | 4A | 19530495 | downstream gene variant | 2011-2012 | -0.257 | -0.090 | 0.236 | $1.03 \times 10^{-4}$ | 0.244 |
| 4328 | 7 | 15043133 | intergenic region | 2011-2012 | 0.084 | 0.021 | 0.236 | $1.22 \times 10^{-4}$ | 0.244 |
| 5755 | 11 | 817810 | intergenic region | 2011-2012 | -0.198 | -0.020 | 0.236 | $7.65 \times 10^{-5}$ | 0.244 |
| 5905 | 11 | 7003007 | splice acceptor variant | 2011-2012 | 0.205 | 0.032 | 0.236 | $1.33 \times 10^{-4}$ | 0.244 |
| 10090 | 24 | 5097562 | upstream gene variant | 2012-2013 | -0.129 | -0.014 | 0.059* | $6.50 \times 10^{-6}$ | 0.072* |

**Table S8:** Autosomal SNPs with significant ($q < 0.25$) net allele frequency shifts between adjacent years in 1999-2013 as detected by [1], reassessed to correct for the larger number of comparisons after inclusion of the Z-linked SNPs and allow for pseudocounts. SNP position is from the Florida Scrub-Jay genome v2, and variant annotation is from SnpEff. $\Delta p$ = change in allele frequency (observed or simulated); 2019 $q$ = adjusted p-value (False Discovery Rate correction) from [1] for multiple comparisons across autosomal SNPs (n = 10,731); $p$ = updated p-value with pseudocounts included; $q$ = adjusted p-value for multiple comparisons across all SNPs in the genome (n = 11,000). SNPs that are significant at a more stringent FDR cutoff of $q < 0.1$ are marked with *.

### Supplementary Figures

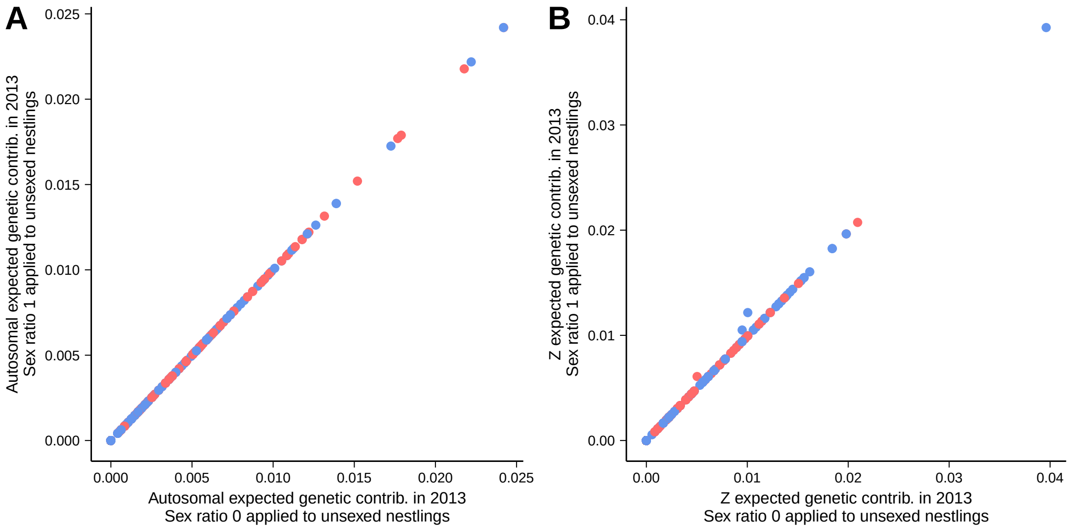


**Figure S1:** Expected genetic contributions to the population in 2013 at (A) a neutral autosomal and (B) a neutral Z chromosome locus for for 351 male breeders (blue) and 378 female breeders (red) estimated by assigning all unsexed individuals as females (sex ratio 0) or males (sex ratio 1).

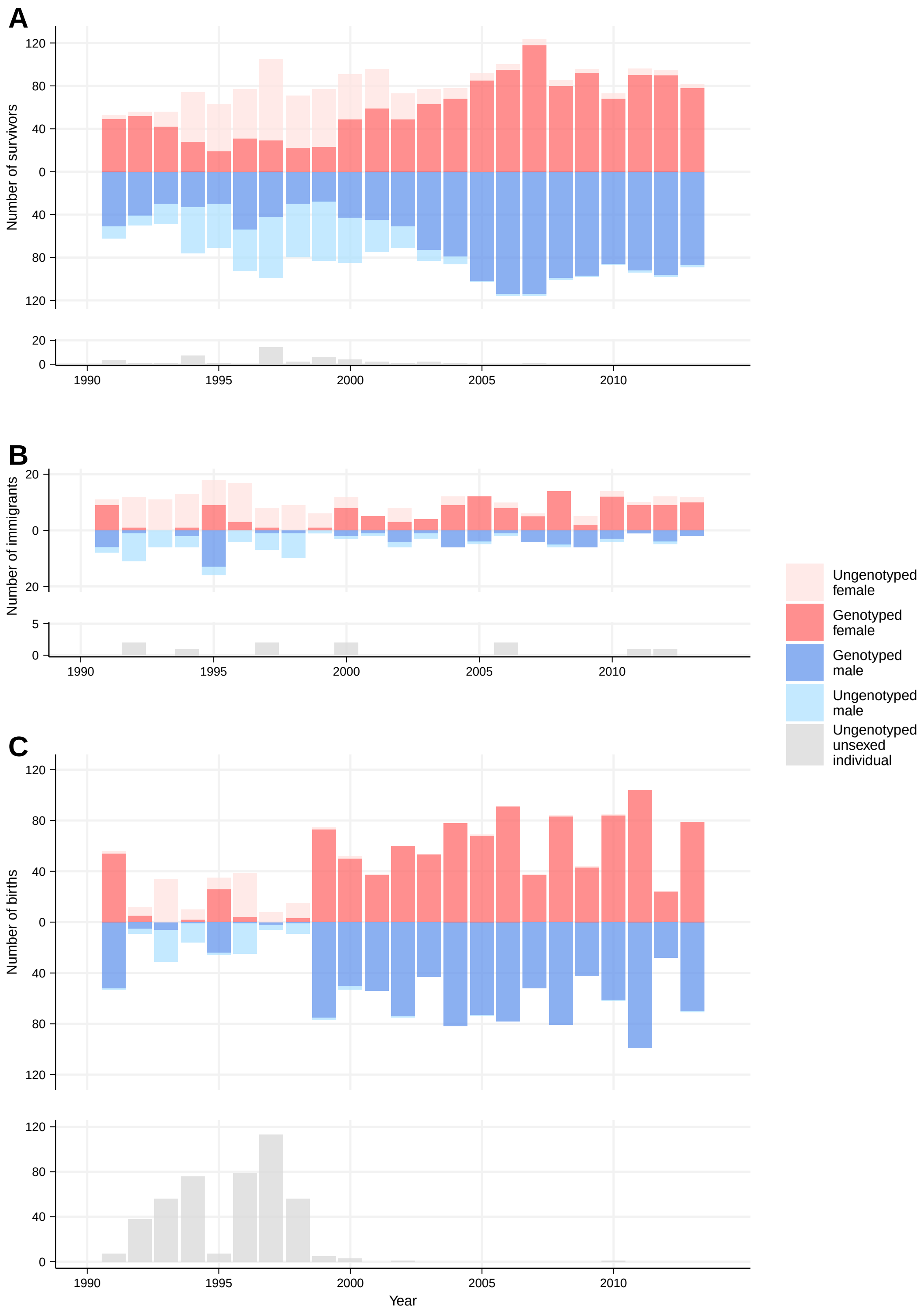


**Figure S2:** Number of sexed and unsexed, genotyped and ungenotyped birds included in each category of the allele frequency model each year: (A) survivors, (B) immigrants, and (C) nestlings (births). Females are shown in shades of red, males are shown in shades of blue, and unsexed individuals are shown in gray.

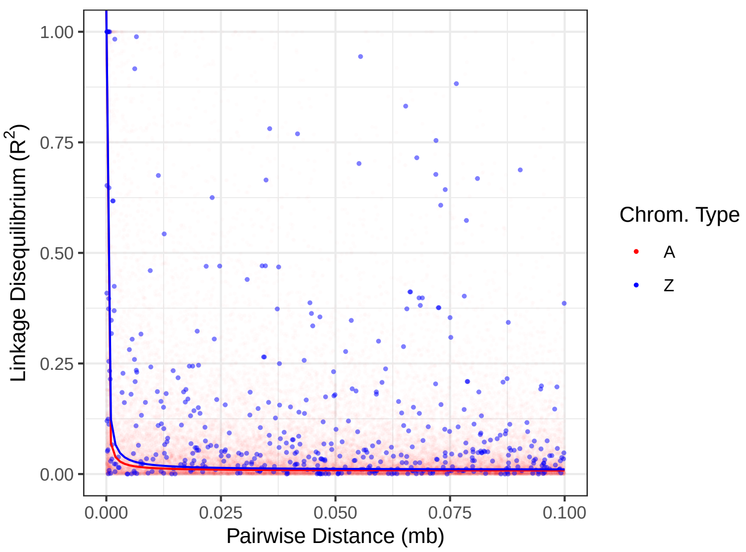


**Figure S3:** Estimated pairwise linkage disequilibrium (LD) between SNPs in each linkage group. Solid lines show the fitted exponential decay curve for the relationship between pairwise LD and distance between SNPs for autosomes in red and the Z chromosome in blue.

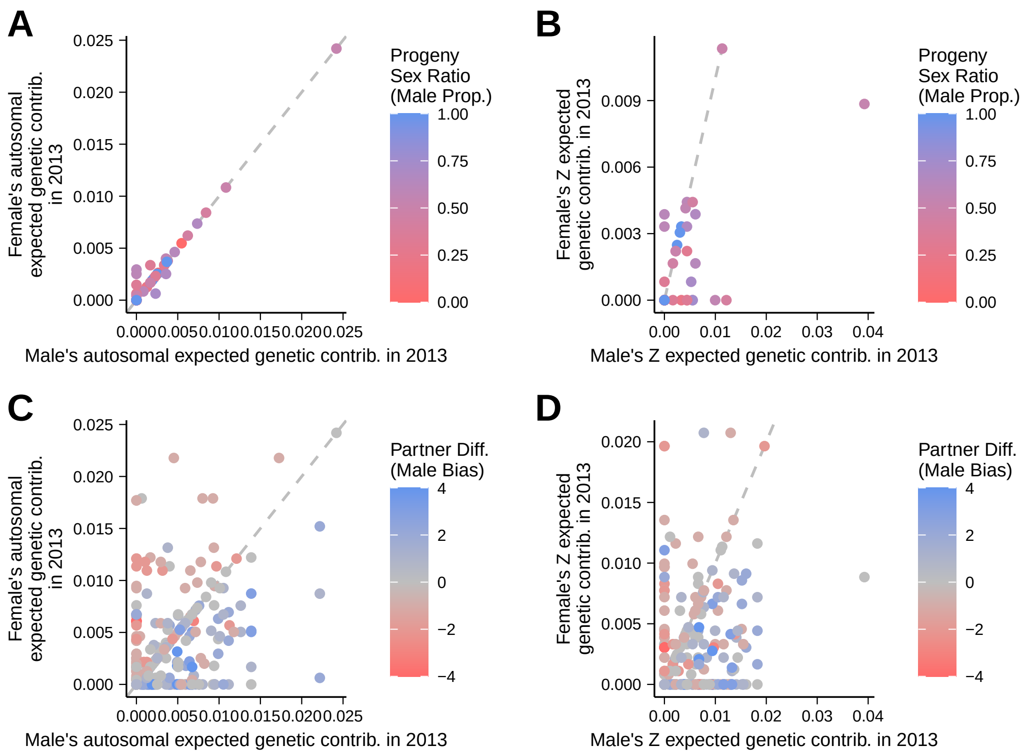


**Figure S4:** Expected genetic contributions between mates for individuals that (A, B) only paired with each other or (C, D) paired with multiple individuals throughout their lifetime on autosomal (A, C) and Z-linked (B, D) loci. The dotted line in each panel shows the 1:1 relationship. In panels A and B, color indicates the progeny sex ratio (blue means more male-biased), while in C and D, color shows the difference in the number of mates each individual had during their lifetime (blue means the male had more mates than the female).

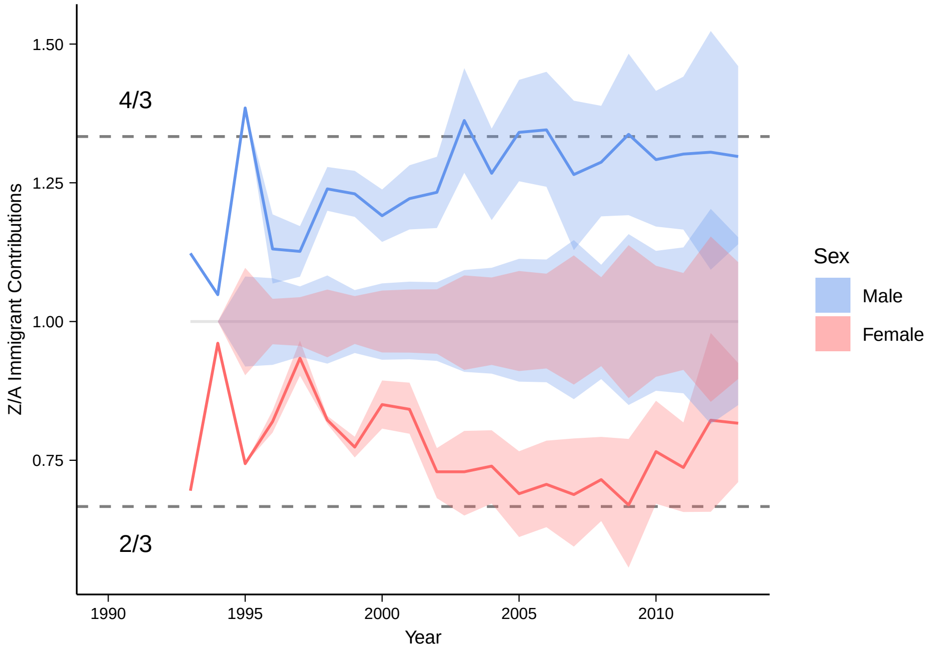


**Figure S5:** Ratio of Z to autosomal expected genetic contributions of male (blue) and female (red) immigrants. The ratio of autosomal to autosomal expected genetic contributions (which should be 1:1) is shown in the center in overlaid blue and red. Dashed lines show the expected ratios. Ribbons show the 95% confidence intervals from the simulations.

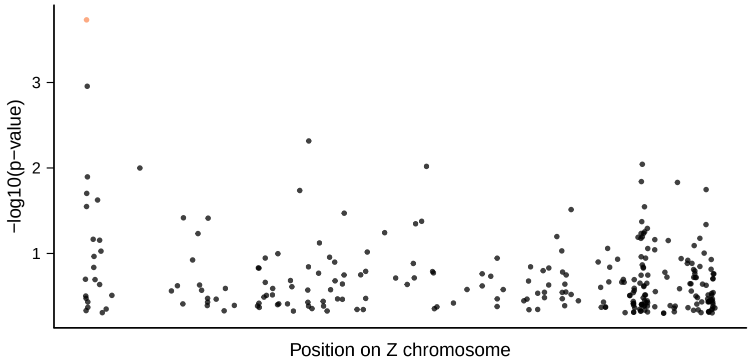


**Figure S6:** Manhattan plot showing allele frequency shifts in 1999-2013. Significant SNPs (under a lenient FDR $< 0.25$) are marked in orange.

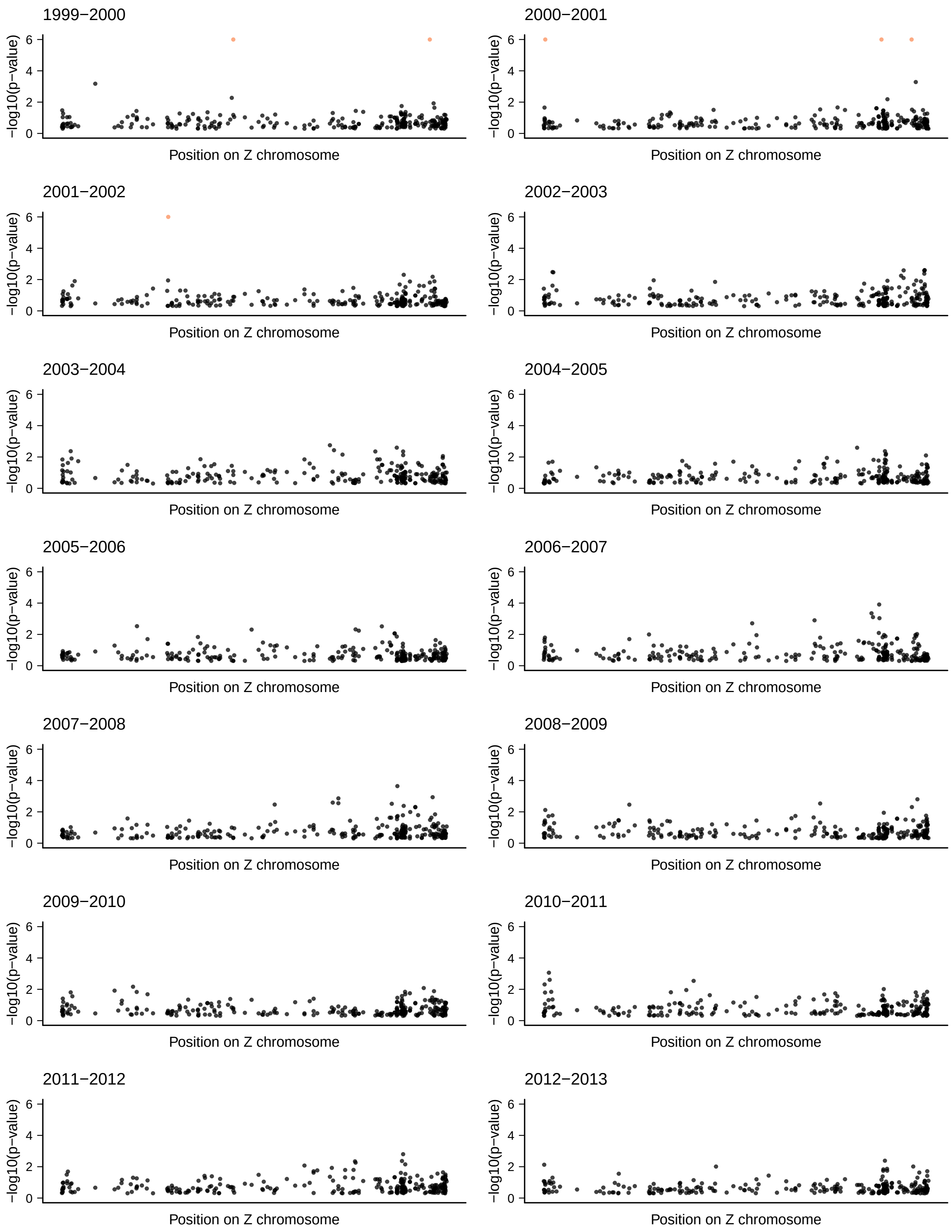


**Figure S7:** Manhattan plot showing allele frequency shifts between every pair of consecutive years in 1999-2013. Significant SNPs (FDR $< 0.1$) are marked in orange.

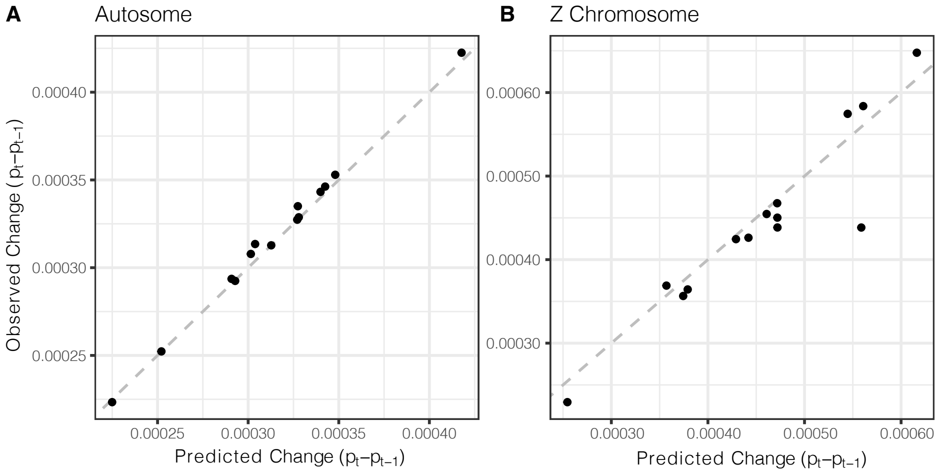


**Figure S8:** Comparison of predicted change in allele frequency across years and the expected change when comparing allele frequency between years on (A) autosome and (B) Z chromosomes. Dashed line shows 1:1.

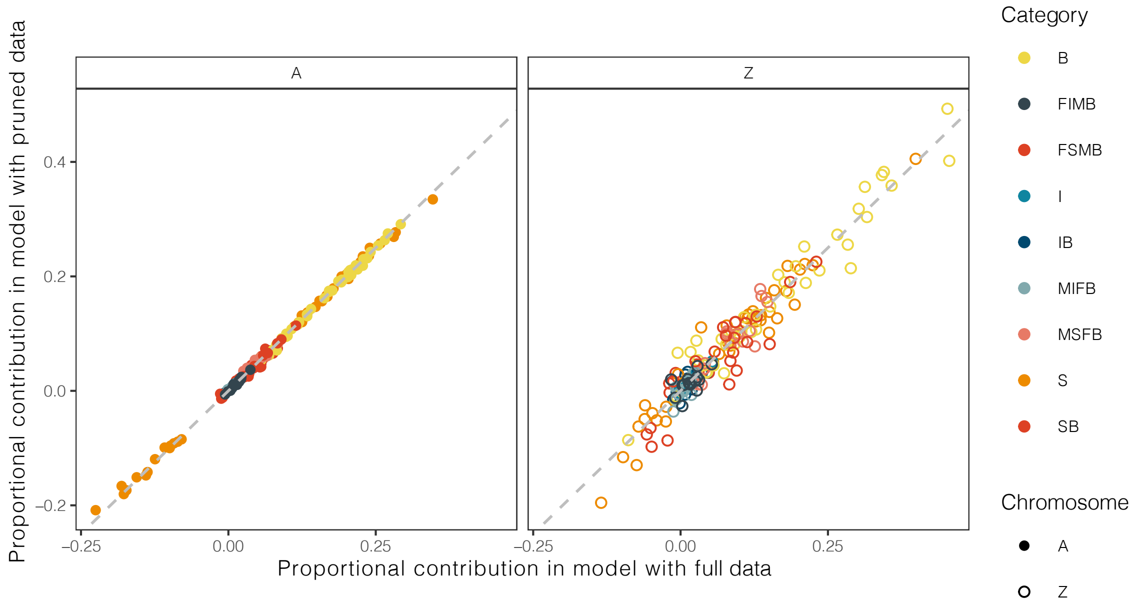


**Figure S9:** Comparison of allele frequency partitioning results for dataset including all SNPs and dataset including and LD-pruned dataset for each category (color) on autosome (solid circle; left panel) and Z (hollow circle; right panel). Dashed line shows 1:1.

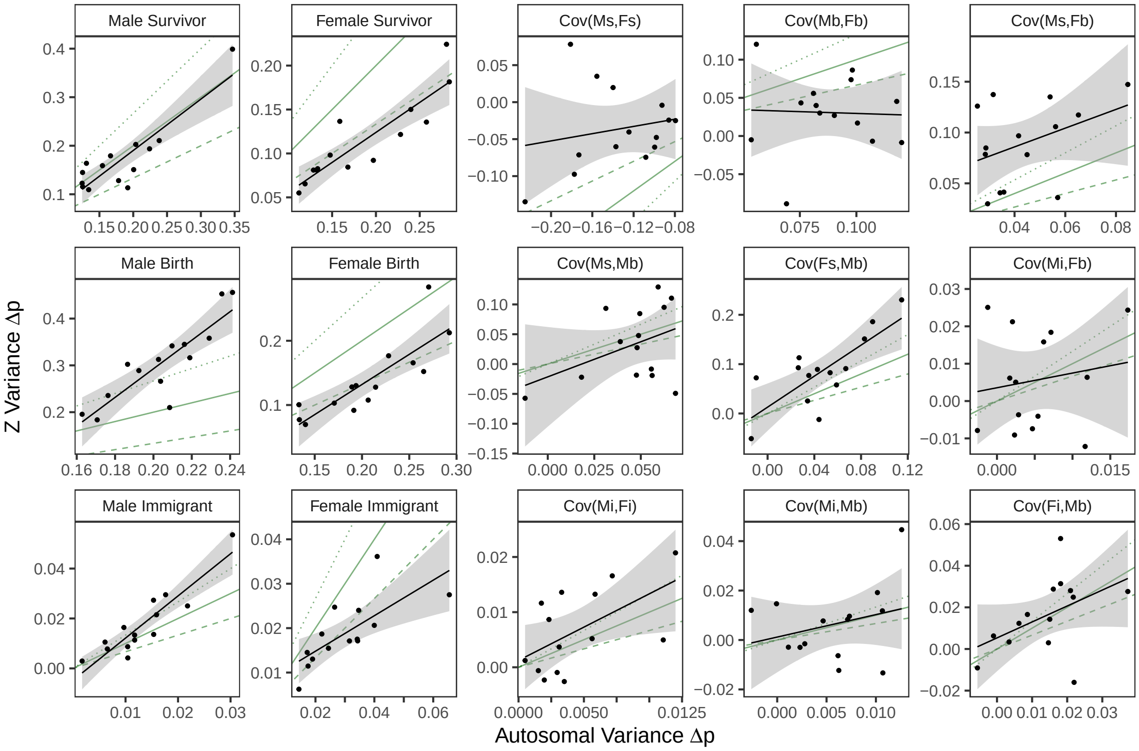


**Figure S10:** Comparison of allele frequency partitioning model results for Z-linked vs autosomal loci. Each panel shows the relationship between Z chromosome allele frequency variance (on the y-axis) and autosomal allele frequency variance (on the x-axis) for the given variance or covariance term. Points show different years. Black lines and grey ribbons show linear models and associated standard error. Solid green lines show a 1:1 relationship, dashed green lines show a 2:3 relationship, and dotted green lines show a 4:3 relationship. Ms = male survivor, Mb = male birth, Mi = male immigrant, Fs = female survivor, Fb = female birth, and Fi = female immigrant.

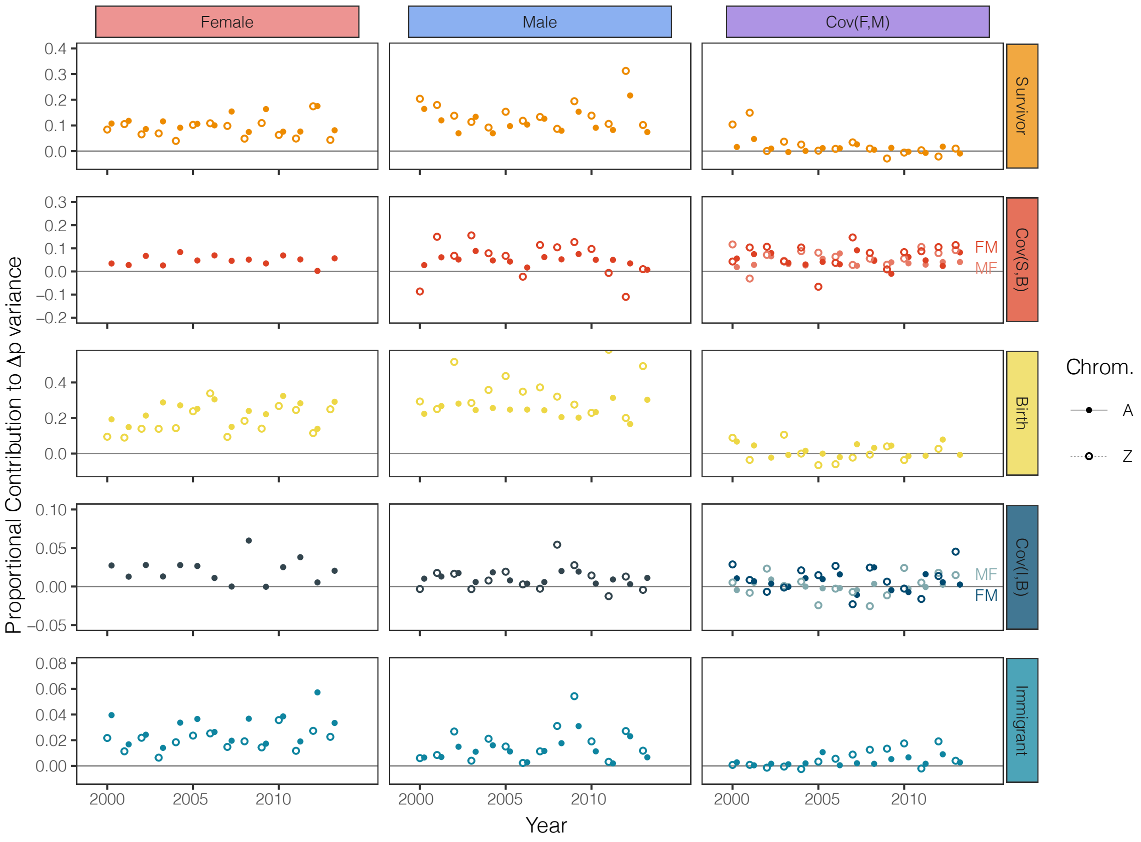


**Figure S11:** Variance contributions of three demographic processes among the sexes across years in a model that accounts for sex-specific allele frequencies in the previous year $(t-1)$. Solid circles and solid lines show the estimates for autosomal loci; open circles and dotted lines show the estimates for Z-linked loci. Cov(S,B) is the covariance between survivors and births, and Cov(I,B) is the covariance between immigrants and births. In the plot showing the covariance between survivors and births of different sexes, the lines labeled "MF" indicate cov(male survivors, female births), and the lines labeled "FM" indicate cov(female survivors, male births); likewise for the plot of the covariance between immigrants and births of different sexes. Note that the y-axis range varies among rows.

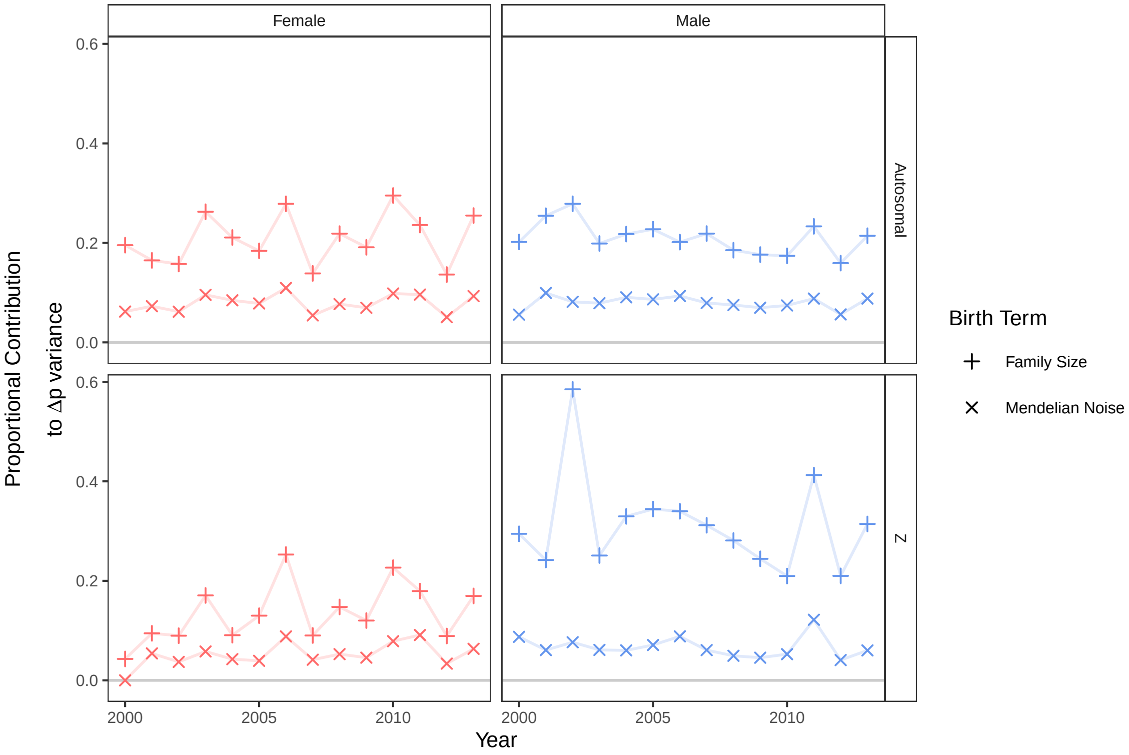


**Figure S12:** Variance in allele frequency change due to variation in family size (+) and Mendelian noise (x) for each sex (female in red, male in blue) across years for autosomal and Z-linked loci.

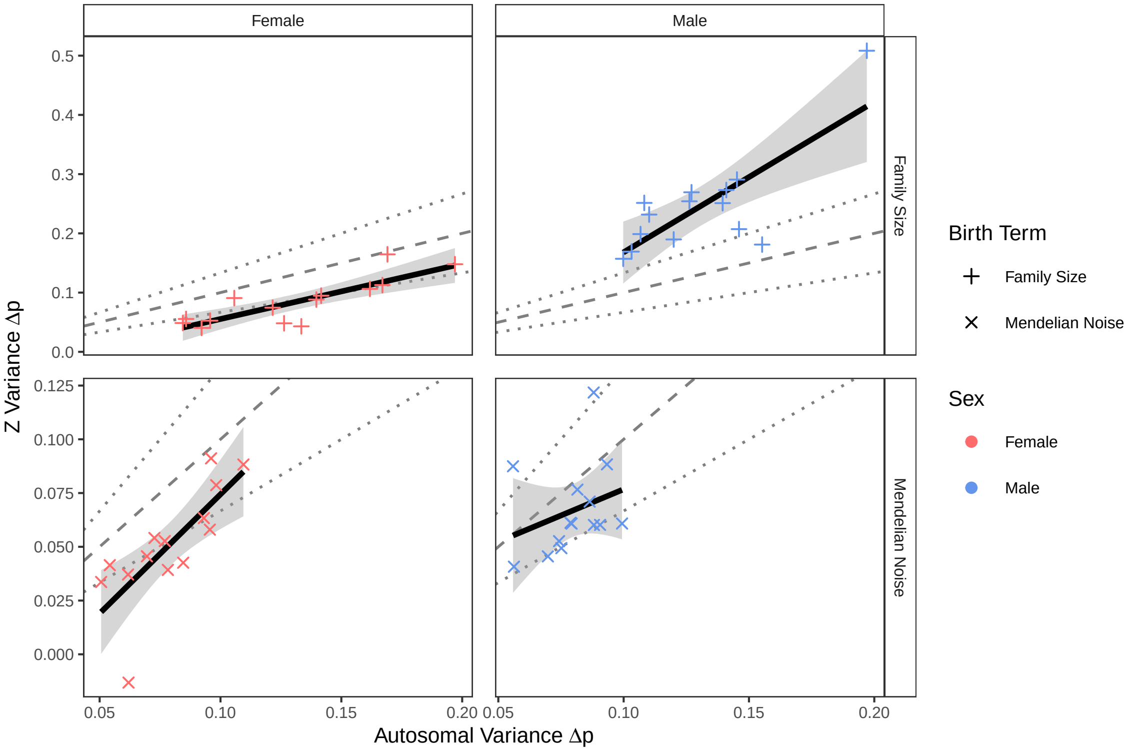


**Figure S13:** Comparison of the variance in allele frequency change due to variation in family size (+) and Mendelian noise (x) for each sex for Z-linked loci vs autosomal loci. Each panel shows the relationship between Z chromosome allele frequency variance (on the y-axis) and autosomal allele frequency variance (on the x-axis) due to variation in family size (top) and Mendelian noise (bottom) for females (left) and males (right). Points show different years. Black lines and grey ribbons show linear models and associated standard error. Dashed gray lines show a 1:1 relationship, and dotted gray lines show a 2:3 or 4:3 relationship.
